# Heat-induced increases in xylem vulnerability: Experimental evidence and possible mechanisms

**DOI:** 10.64898/2026.11.05.764619

**Authors:** Roman Mathias Link, Sharath Shyamappa Paligi, Namalu Abeysundara, Christian Ammer, Janithya Dishani Kodituwakku Dharmawardhane, Kerstin Pierick, Abdul Razzak, Bernhard Schuldt, Fon Robinson Tezeh, William Robert Wighton, Kate M Johnson, Ilaine Silveira Matos

**Author notes:** Shared first authorship:* Roman Mathias Link & Sharath Shyamappa Paligi. Shared last authorship:* Kate M Johnson & Ilaine Silveira Matos.

## Abstract

- Hotter droughts driven by global warming are increasing tree mortality worldwide. Here, we propose a theoretical framework that explains how dissolved gas affects the stability of bubbles in xylem sap, and shows that in the presence of gas, sudden increases in temperature can accelerate embolism formation. We then provide experimental evidence that heat exposure can change xylem vulnerability to drought.
- Using data from seven Australian and European tree species, we assessed how elevated temperature (+15 and +20°C above ambient) affects both xylem embolism resistance, measured with the Optical Vulnerability Technique, and VPD-corrected drying times.
- We found evidence for reduced embolism resistance and accelerated dehydration in heat-exposed plants. On average, trees reached the water potential at 50% loss of xylem function at 0.73 MPa less negative water potentials and 7.8 kPa h earlier, though responses were highly species-specific, and credibly different from zero only in the Australian species.
- Our results are consistent with the proposed framework and suggest that embolism resistance may not be a static trait, which has important repercussions for studies of plant vulnerability to drought. Further studies are necessary to assess the potential effects of heat-induced reductions in embolism resistance on tree mortality during heatwaves.

## Introduction

Over the last decades, climate-change driven drought events have become a major driver of tree mortality worldwide (Allen et al., 2015; Hammond et al., 2022). Particularly extreme tree mortality events have been linked to global change-type and thus hotter drought conditions that combine reduced soil water availability with extreme atmospheric temperatures and dryness during heatwaves in Europe (Schuldt et al., 2020), Australia (Mitchell et al. 2014; Matusick et al., 2018; Marchin et al., 2022), and across the world (Allen et al., 2015; Breshears et al., 2013). Under such conditions, a disruption of the plant water transport system due to the formation of air emboli has been identified as an important cause of tree death (Arend et al., 2021; McDowell et al., 2022).

How exactly xylem embolism develops under drought – and particularly under combined drought and heat stress – is complex and subject to debate (Lens et al., 2022). The water transport from roots to leaves via the cohesion-tension mechanism (Askenasy, 1895; Dixon & Joly, 1895) requires the liquid in the xylem to be under – often substantial – negative pressures. Under these conditions, water is in a metastable state, and a spontaneous phase transition into the gas phase is theoretically possible. However, the water potentials required to induce homogeneous cavitation, i.e., the spontaneous nucleation of water vapour bubbles under sufficiently negative water potentials, are orders of magnitude more negative than the water potentials usually observed in the xylem (Herbert & Caupin, 2005; Stroock et al., 2014; Kanduc et al., 2020). Instead, it is widely accepted that the major driver of embolism formation is the spread of gas from already embolized to adjacent conduits via pit membranes, although the process driving this embolism propagation is debated (Jansen et al., 2026). The classical explanation for embolism propagation is a process called air seeding, which involves the pressure-driven bulk flow of gas bubbles through pit membrane pores and their subsequent expansion (Lewis 1988; Tyree & Sperry, 1988). Explanations for the initial source of air for air seeding include mechanical damage to the plant (Johnson et al., 2022), and air in cells connected to the xylem, such as imperforate tracheary elements, which may embolise earlier than the xylem conduits (Johnson et al., 2023). As the air-seeding pressure of a pit membrane pore is a function of its diameter, the embolism risk of a conduit would be determined by the size of its largest membrane pore (Wheeler et al., 2005; Hacke et al., 2006; Christman et al., 2009). However, newer evidence on the ultrastructure of pit membranes indicates that large enough pit membrane pores are unlikely to be as common as required for this mode of embolism propagation (Kaack et al., 2021). An alternative explanation for embolism propagation is that air instead enters into adjacent conduits in the form of stable nanobubbles that snap off in the pore constrictions of pit membranes (Schenk et al., 2015). These nanobubbles are then coated by surfactants (Schenk et al., 2017), which stabilize them against expansion due to the resulting dynamic surface tension (Yang et al., 2021; Ingram et al., 2021). Embolism formation happens at a later stage when nanobubbles lose their stability and expand, and is thus temporally and spatially separated from bubble formation (Jansen et al., 2026). Embolism spread may further be aided by gas diffusion from adjacent conduits (Guan et al.; 2021; Pereira et al., 2023). Cavitation events in the true sense of the word (i.e., spontaneous bubble expansion rather than the growth of existing gas-filled cavities) are likely rare. Mechanistic modelling suggests that they are most likely to happen in lipid bilayers present in the sap, and unlikely to be quantitatively relevant at water potentials less negative than −10 MPa (Kanduc et al., 2021).

It has long been known that xylem sap can contain high concentrations of dissolved gas (see e.g. Scholander et al., 1955). Plants take up water from the soil, which is a complex porous interface where – unless fully saturated – water is always in direct contact with air over a large surface area. It can therefore be assumed that soil water is more or less gas-saturated (Marion et al., 2026). Inside the xylem, dissolved gas is in equilibrium with the gas phase in air-filled spaces, e.g. in fibres, embolised conduits and intercellular spaces in the mesophyll. As gas solubility is extremely temperature-dependent, with solubility decreasing with increasing temperature (Box 1), conditions of supersaturation have long been assumed to happen regularly driven by diurnal temperature cycles (Hammel et al., 1967; Sperry & Sullivan, 1992; Cobb et al., 2007), as demonstrated experimentally by Schenk et al. (2016). Nonetheless, in a study of the molecular dynamics and kinetics of cavitation processes in the xylem, Kanduc et al. (2020) concluded that dissolved gases at low concentrations do not significantly modify the time to cavitation events compared to pure water. However, recent analyses of the thermodynamics of bubble nucleation on preexisting nuclei indicate that gas content can in fact significantly reduce the energy barrier for bubble nucleation compared to bubbles consisting of pure water vapour, and completely eliminate it if a bubble is supersaturated with gas compared to the equilibrium state (Alamé & Mahesh, 2024). Spontaneous bubble nucleation in the xylem from gas-saturated nuclei might thus require less extreme negative pressures than previously expected. Consequently, the presence of dissolved gas in the xylem may have a strong impact on embolism formation.

The solubility of the four most common gases in water declines by between −52.1 and −72.8% over the biologically relevant temperature range from 0 to 50 °C (Fig. 1a, Box 1). Over the same range, surface tension and viscosity, two other highly relevant properties for bubble nucleation and hydraulic processes in the xylem, decline by −10.2 and −40.3%, respectively (Box 1, Fig 1b, 1c). The temperature rise on extremely hot days may push xylem sap into conditions of strong gas supersaturation, and potentially induce bubble nucleation at less negative water potentials compared to non-heated conditions. The same process can also result in embolism formation in blood vessels in the human body even in the absence of negative pressures, e.g. when hypothermic patients are rewarmed too rapidly (Geissler et al, 1997; Mardanpour, 2026), when hyperthermic intraperitoneal chemotherapy is administered (Vukadinovic et al., 2015; Mardanpour, 2026), or when blood transfusions are saturated at low temperatures and rewarmed rapidly prior to infusion (Mashimo, 1980). If a similar temperature-driven outgassing happened in plants, it would be expected to push vulnerability curves (i.e. the relationship between cumulative embolism and water potentials) towards less negative water potentials.

**Figure 1.**
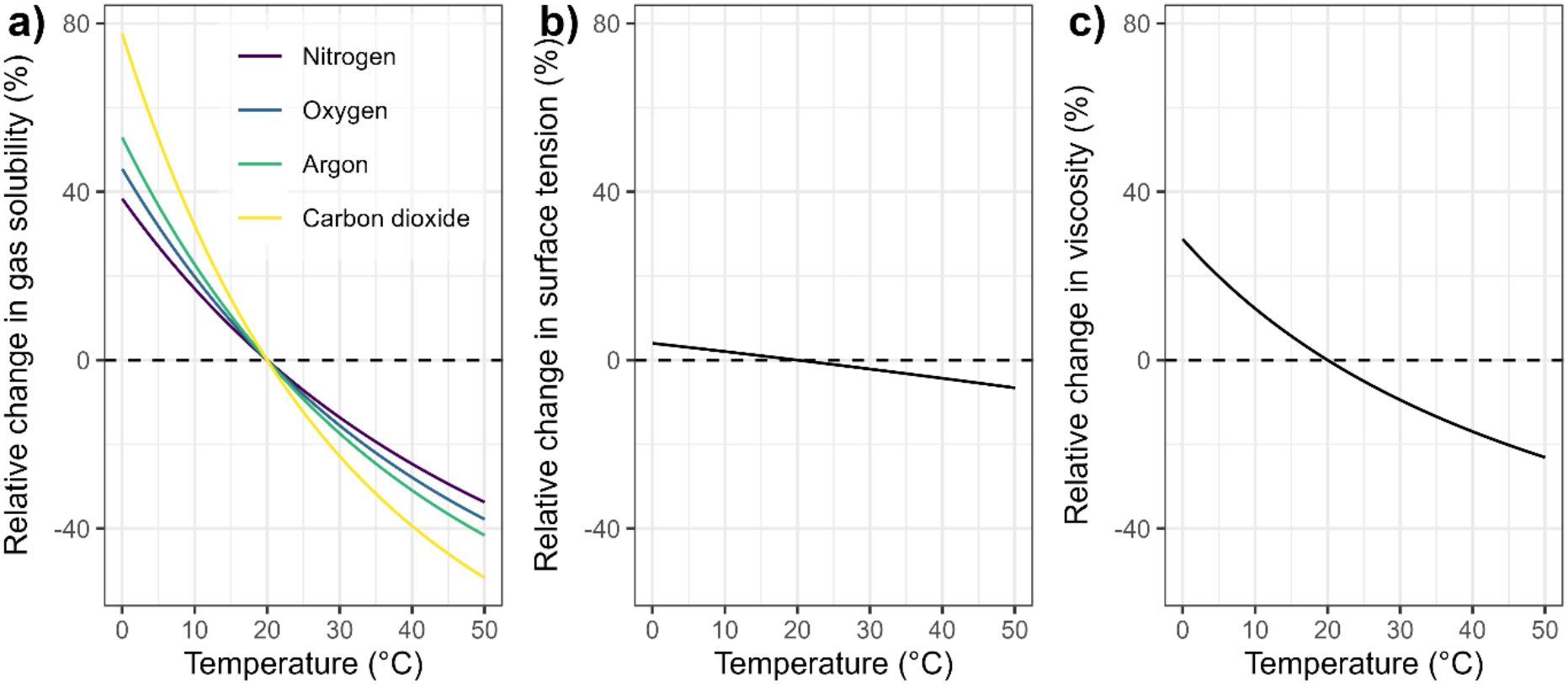
Relative change in a) gas solubility, b) surface tension and c) viscosity of pure water over the temperature range from 0 to 50 °C compared to their reference value at 20 °C. In a), the change in solubility is shown for the 4 most common gases in the atmosphere besides water vapour. The values were calculated according to the equations in Box 1, using values from Sander et al. (2023) for the computation of gas solubility.

To our knowledge, so far, there are few studies on temperature-dependent changes in vulnerability curves. An increased vulnerability to embolism at higher temperatures was demonstrated experimentally by Cochard et al. (2007). Further evidence comes from studies of fire effects, which suggest a reduced embolism resistance (Michaletz et al, 2012; Lodge et al., 2018; Bär et al., 2018) and an increase in native embolism (West et al., 2016) after heat exposure in water baths or simulated heat plumes, which these authors at least partially attribute to declining air-seeding pressures due to the reduced surface tension at higher temperatures. For the same reason, Cochard (2021) predicted a shift in the water potential at 50% loss of conductivity by about +0.2% per °C (Fig. 1c). We argue that the magnitude of the changes in surface tension alone is likely not sufficient to explain observed shifts in vulnerability curves, and propose that the strong temperature sensitivity of gas solubility (Fig. 1a) paired with the effect of dissolved gases on bubble nucleation warrant further investigation.

Here, we present a theoretical framework for bubble nucleation in the presence of dissolved gases based on Alamé and Mahesh (2024), and highlight how rapid temperature surges may drive embolism formation by destabilizing existing nuclei. We find experimental evidence for increased vulnerability to xylem embolism under elevated temperature, providing potential support for this mechanism. Through two experiments, conducted in Australia and Germany, we investigated whether leaf embolism formation is temperature dependent. In both experiments, tree saplings were subjected to heat treatments in climate chambers, and leaf vulnerability curves obtained with the Optical Vulnerability Technique (Brodribb et al., 2016) were compared between controls and heat treatments. In the German branch of the experiment, we further experimentally supersaturated the water in the xylem with air before dehydration. We used these two datasets to test the hypotheses that (i) the water potential at 12, 50 and 88% cumulative embolism (*P*_12_, *P*_50_ and *P*_88_, MPa), as well as the expected value of the embolism pressure distribution (*P*_ex_, MPa), are shifted towards less negative values under hotter-drought conditions, (ii) after correcting for differences in vapour pressure deficit at different temperatures, the time to reach 12, 50 and 88% cumulative embolism (*T*_12_, *T*_50_ and *T*_88_, kPa h), as well as the expected value of the drying time distribution (*T*_exp_, kPa h), are shorter in the heat treatment.

### Description

#### Bubble nucleation in pure water

Following the framework of classical nucleation theory (Sear et al., 2007), the Gibbs free energy *ΔG*_*tot,v*_ of a spherical cavity in pure water can be described as:

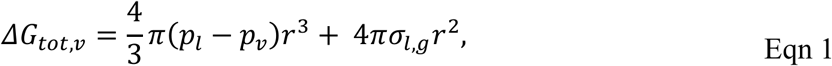

where the indices *v, l* and *g* denote the vapour, liquid and gas phase, respectively, *r* is the bubble radius (m), *p*_*v*_ is the vapour pressure inside the gas bubble (Pa), *p*_*l*_ is the liquid pressure (Pa) and *σ*_*l,g*_ is the surface tension at the liquid-gas interface (J m^−2^). The first term describes the work performed by the volume expansion, while the second term describes the free energy required for creating the bubble interface (Kanduc et al., 2021).

By setting the first derivative of Eqn 1 with respect to *r* to zero and rearranging, an equilibrium solution can be found which results in the critical radius *r*_*cr,v*_:

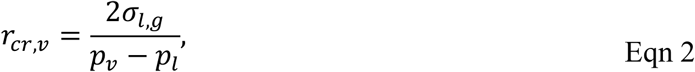

The energy barrier 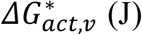 required to surpass the critical radius *r*_*cr,v*_ follows analogously as:

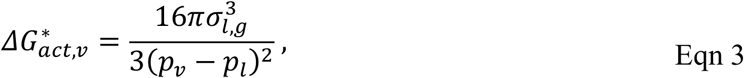

#### Bubble nucleation in the presence of dissolved gas

In the presence of gas, a free bubble is stable when the expanding forces (i.e., pressure of the gas in the bubble and vapor pressure) are equal to the collapsing forces (i.e., pressure of the liquid and effects of surface tension):

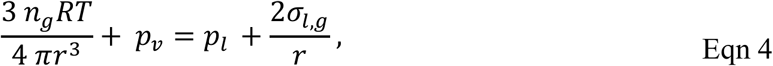

where *n*_*g*_ is the mole number of the gas in the bubble (mol), *p*_*v*_ is the vapour pressure (Pa), *R* is the gas constant (8.3145 J K^−1^ mol^−1^) and *T* is the temperature in K.

For a bubble radius to be stable, the rate of change of the expanding and collapsing forces has to be equal. Taking the derivative of both sides of the equation and rearranging results in the following expression for the critical radius in dependence of gas content:

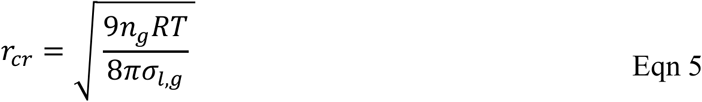

Inserting this value for *r*_*cr*_ into the Laplace equation results in the following equation for the critical radius, which is known as Blake’s radius *r*_*cr,Bl*_ (Blake 1949a,b):

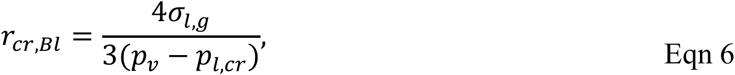

where *p*_*l,cr*_ (Pa) is the critical pressure.

While this is a valid description for the critical radius under equilibrium conditions, Alamé and Mahesh (2024) point out that in itself it is not a complete cavitation model. These authors instead used a Gibbs free energy approach to derive a more accurate expression of the free energy of a bubble that takes into account the chemical equilibrium between the gas phase in the bubble and the surrounding liquid (for a full derivation, see Alamé and Mahesh, 2024).

According to their framework (upon which the rest of this section is based on), the free energy *ΔG*_*tot,v*+*g*_ (J) of a spherical bubble in water containing dissolved gases is as follows:

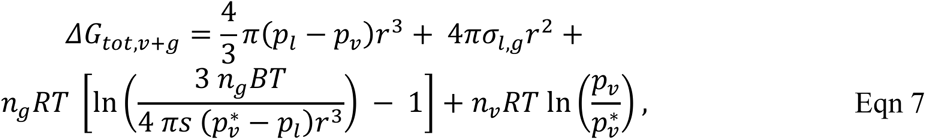

where *n*_*g*_ and *n*_*v*_ are the mole number of the gas and water vapour in the bubble (mol), *p*_*v*_ is the current and 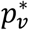 the saturation vapour pressure in the bubble (Pa), *R* is the gas constant (8.3145 J K^−1^ mol^−1^), *T* is the temperature in K, and *s* is the degree of gas saturation expressed as the ratio of the current mole fraction of dissolved gas (*x*_*dg*_) to the saturated one 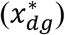, i.e. 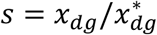 (unitless). Here, as in Eqn 1, the first and second term describe the work performed by the volume expansion and the free energy required for creating the bubble interface. The third and fourth terms together describe the total chemical potential energy, with the third term describing the difference in chemical potential between the free gas in the bubble and the dissolved gas in the liquid phase, and the fourth term describing the potential difference between unsaturated and saturated vapor phase. Fig. 2a shows the relationship between *ΔG*_*tot,v*+*g*_ and bubble radius for different gas contents under the incipient regime (*p*_*l*_ − *p*_*v*_ < 0), with a visible decline in the nucleation barrier for higher degrees of gas saturation. An analogous plot for the desinent regime (*p*_*l*_ − *p*_*v*_ > 0) is shown in Fig. S1.

**Figure 2.**
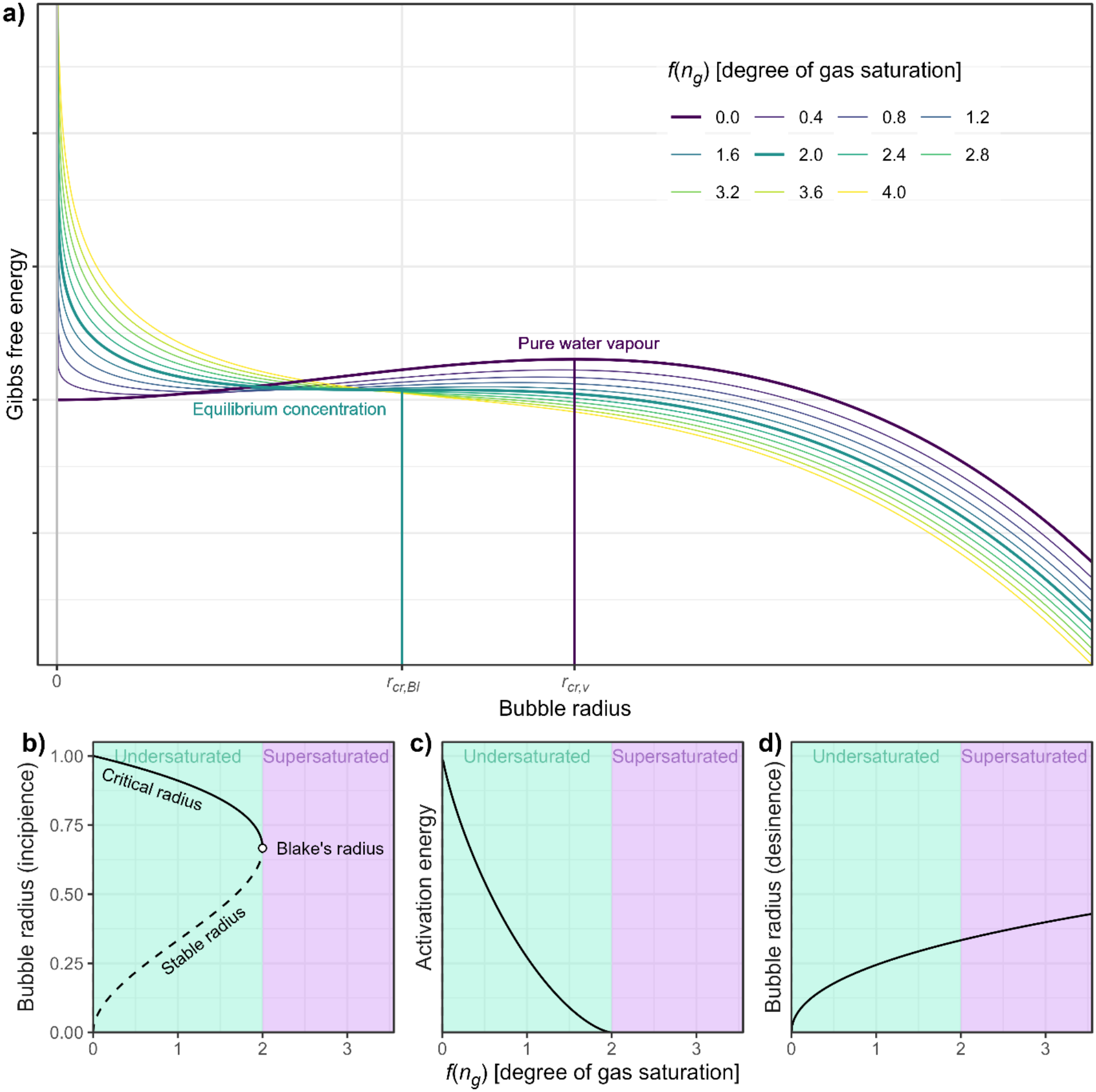
a) Change in Gibbs free energy with bubble radius for different gas contents *n*_*g*_ under the incipient regime. Shown are the (relative) values of *ΔG*_*tot.v*+*g*_ following Eqn 7 coloured by *f*(*n*_*g*_) (an indicator of the degree of gas saturation of the gas in the bubble as a function of gas content, ranging from 0 for pure water vapour over 2 for equilibrium saturation to > 2 for supersaturated conditions; see Eqn 10). Highlighted radii are the critical radius *r*_*cr,v*_ following Eqn 2 for pure water vapour (gas content of 0) and Blake’s radius *r*_*cr,Bl*_ following Eqn 6 for equilibrium conditions. b) Change in the critical radius *r*_*cr,v*+*g*_ for incipient cavitation and the equilibrium radius *r*_*eq*_ with *f*(*n*_*g*_) relative to the critical radius *r*_*cr,v*_ for pure water vapour. For *f*(*n*_*g*_) = 2, both radii are identical and recover Blake’s radius *r*_*cr,Bl*_. For values of *f*(*n*_*g*_) > 2 (supersaturation), no critical radius is defined. c) Relative change in the activation energy needed for bubble expansion 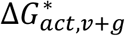 compared to pure water vapour plotted against *f*(*n*_*g*_). For supersaturated conditions, no energy barrier is defined. d) Change in the equilibrium radius *r*_*eq*_ for the desinent regime with *f*(*n*_*g*_) relative to the critical radius *r*_*cr,v*_ for pure water. Here, values are also defined for the supersaturated range: more dissolved gas permits higher stable bubble radii.

A critical radius *r*_*cr,v*+*g*_ (m) for incipient cavitation (*p*_*l*_ − *p*_*v*_ < 0) can be derived by setting the first derivative of Eqn 7 with respect to *r* and *n*_*g*_ to zero and solving the resulting cubic equation (Alamé and Mahesh, 2024). The solution can be expressed as the product of the critical radius *r*_*cr,v*_ in pure water and a correction factor *ξ*_*inc*_(*n*_*g*_) depending on gas content:

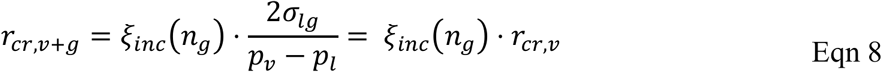

The correction factor *ξ*_*inc*_(*n*_*g*_) is as follows:

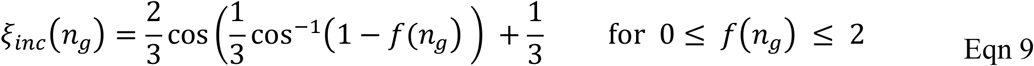

Here, *f*(*n*_*g*_) is an equation that describes the effect of gas content on the stability of the bubble:

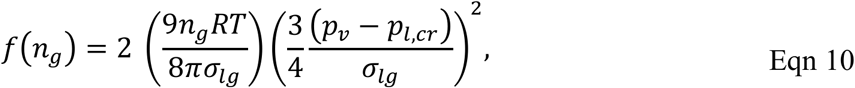

For pure water vapour, *n*_*g*_ is 0, resulting in a value of 0 for *f*(*n*_*g*_) and hence a value of 1 for *ξ*_*inc*_(*n*_*g*_), recovering the expression for *r*_*cr,v*_ from Eqn 2 when inserting into Eqn 8. If the expanding and contracting forces in the bubble are in equilibrium, the following equality holds (compare Eqn 5, 6):

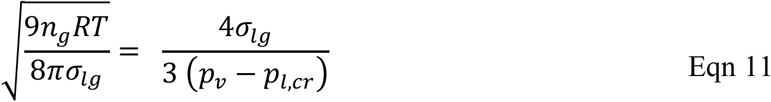

Accordingly, when the bubble is in equilibrium, *f*(*n*_*g*_) is equal to 2, resulting in *ξ*_*inc*_(*n*_*g*_) = 2/3, in which case *r*_*cr,v*+*g*_ evaluates to Blake’s radius *r*_*cr,Bl*_ from Eqn 6. For values of 0 < *f*(*n*_*g*_) < 2, the bubble is undersaturated and for values of *f*(*n*_*g*_) > 2 supersaturated with gas compared to equilibrium conditions (Fig. 2b). Note that Eqn 9 is only defined for *f*(*n*_*g*_) ≤ 2, reflecting that no critical radius exists if the bubble is supersaturated, as the activation barrier required for nucleation is then reduced to 0 (see below).

A stable radius *r*_*eq*_ (m) for incipient cavitation (*p*_*l*_ − *p*_*v*_ < 0) can be derived analogously:

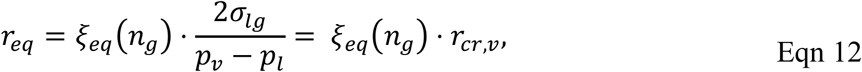

With the correction following correction factor *ξ*_*eq*_(*n*_*g*_):

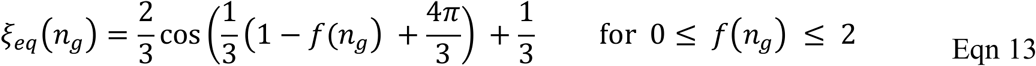

The equilibrium concentration is 0 for pure water vapour and rises with *f*(*n*_*g*_) until reaching the same value as *r*_*cr,v*+*g*_ at *f*(*n*_*g*_) = 2, again resolving to Blake’s radius (Fig. 2b). This can be seen in Fig. 2a, where *r*_*eq*_ corresponds to a local minimum in free energy for 0 < *f*(*n*_*g*_) < 2. For *f*(*n*_*g*_) = 2, the minimum disappears, and the free energy has a saddle point at Blake’s radius.

The energy barrier 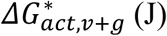 required to surpass the critical radius *r*_*cr,v*+*g*_ can also be solved analytically (Alamé and Mahesh, 2024):

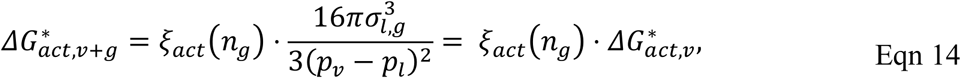

where *ξ*_*act*_(*n*_*g*_) is a correction factor for the effect of gas content that is defined as follows:

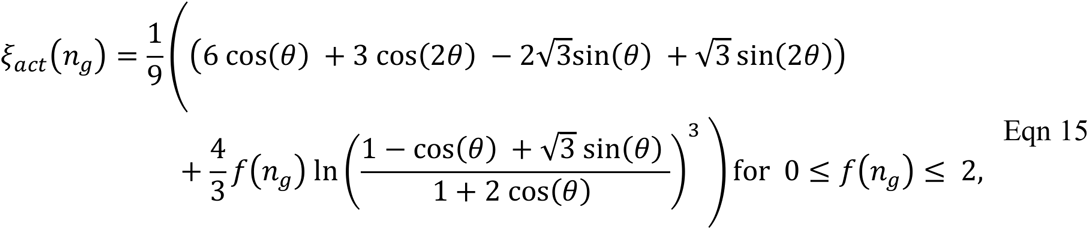

where 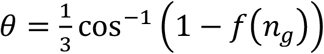. Accordingly, the energy barrier 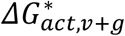 is equal to the value of 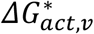 for pure water vapour from Eqn 3 and declines with increasing gas content, reducing to zero for equilibrium conditions (*f*(*n*_*g*_) = 2). For supersaturated conditions, 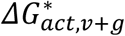 is not defined (Fig. 2b).

For desinent conditions when negative pressure subsides (*p*_*l*_ − *p*_*v*_ > 0), Alamé and Mahesh (2024) provide a solution for a new equilibrium radius *r*_*eq,des*_ (m):

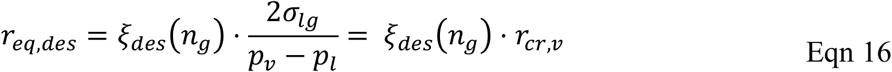

Here, the correction factor *ξ*_*des*_(*n*_*g*_) for the effect of gas content is defined as follows:

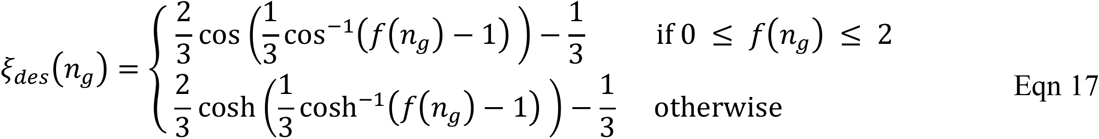

Accordingly, for pure water vapour, the equilibrium radius under a desinent regime is equal to 0, and increases with higher gas concentrations into the supersaturated range (Fig. 2d, Fig. S3). Hence, the presence of gas prevents complete bubble dissolution, and the more moles of gas a bubble contains, the higher its stable equilibrium radius.

#### Implications for xylem embolism formation

The model of bubble nucleation by Alamé and Mahesh (2024) has several important implications for the study of xylem embolism formation, most importantly, a) in a bubble with gas content under equilibrium conditions, the energy barrier for expansion is effectively reduced to zero (Fig. 2c), b) the critical radius for bubble expansion is highest for pure water vapour bubbles, decreases with gas content and is not defined for bubbles in supersaturated conditions, as under those conditions there is no energy barrier for expansion (Fig. 2b), and c) if they contain gas, bubbles below the critical radius shrink to their stable equilibrium radius *r*_*eq*_ rather than dissolving completely (Fig. 2b), even after the negative pressure subsides (Fig. 2d, Fig. S1).

While it is possible that bubbles with a non-zero gas content form and expand spontaneously, even in the presence of considerable dissolved gas it is much more likely that expansion happens from pre-existing nucleation sites. Potential nucleation sites in xylem sap are hydrophobic surfaces, surfactant micelles or lipid bilayers (Kanduc et al., 2021), and, importantly, stable nanobubbles present in the sap (Schenk et al., 2015, 2017). While the spontaneous expansion of nanobubbles is a potential explanation of embolism formation, the factors that cause this expansion are not yet fully resolved (Jansen et al., 2026). High concentrations of dissolved gas are assumed to stabilize nanobubbles against dissolution (Schenk et al., 2015, 2016), which is confirmed by the existence of non-zero stable radii for bubbles below the critical radius derived above.

However, our framework also indicates that existing bubbles may be destabilized when temperature rises. On the short term, a rise in temperature will increase *f*(*n*_*g*_) directly because it scales linearly with temperature and to the third power with inverse surface tension (Eqn 10). In consequence, rapid surges in temperature can cause a change in the equilibrium state that results in previously stable bubbles now containing enough gas to push them over their critical radius. Over longer time scales, higher temperatures can moreover increase the gas content *n*_*g*_ in the bubble itself due to diffusion driven by the reduced gas solubility of the sap (Fig. 1a, Eqn B1.3). Even on most regular days, it is likely that xylem sap goes through periods of slight supersaturation during warm periods (Schenk et al., 2016). During extreme heatwaves, the effect of strong temperature differentials and the resulting highly supersaturated conditions may drive the spontaneous expansion of bubbles and result in xylem embolism.

## Materials and methods

### Plant material

Both experiments were conducted on potted saplings, with a total of eight angiosperm tree species assessed, one of which had to be discarded due to data quality issues.

In Adelaide, South Australia, five *Eucalyptus* tree species (*E. argophloia* Blakely – subtropical to mediterranean climate, *E infera* A.R.Bean – subtropical climate, *E. macrocarpa* (Maiden) Maiden – Mediterranean climate, *E. occidentalis* Endl. – temperate climate, *E. camaldulensis* Dehnh. – broad distribution across Mediterranean, temperate, tropical and sub-tropical climates) were grown from seeds purchased from the Australian Tree Seed Centre or were obtained as tube stocks from local tree nurseries. All saplings were grown in 10 L native pot mix soil (Scotts Osmocote, Bella Vista, NSW, Australia) under well-watered conditions and natural sunlight, and fertilized every six months (Native Scotts Osmocote Controlled Release Fertiliser, Bella Vista, NSW, Australia) to achieve maximum growth. Once saplings were about 1–2 years old (Jan–Feb 2026), five replicates per species per treatment were selected for the xylem vulnerability measurements, prioritising saplings with similar height and healthy-looking leaves. Before the measurements, the pots were watered to full capacity, and the whole saplings were covered with a dark plastic bag and rehydrated overnight to full turgor.

In Göttingen, Germany, we used each 10 saplings of the three temperate broadleaved tree species *Alnus glutinosa* (L.) Gaertn., *Castanea sativa* Mill. and *Prunus avium* L. In 2026, two-year-old saplings were sourced from local tree nurseries and potted into 12 L pots using a soil mixture of medium-textured peat mix and medium grained fine sand, and then grown in full sun in the Experimental Botanical Garden Göttingen until their harvest in July 2026. Before measurement, we bare-rooted the saplings and washed their rootstock to remove all remaining soil substrate to speed up dry-down. We then hydrated the plants overnight at 15 °C in a dark room with water that we saturated with gas by bubbling it with compressed air for at least 2 h to ensure supersaturation of the water in the xylem. After hydration, we towel-dried their rootstock and moved them to the climate chambers where the dry-down experiment took place.

### Image acquisition

Vulnerability to xylem embolism was assessed in all samples using the Optical Vulnerability Technique (OVT; Brodribb et al., 2016).

*Australia:* saplings were bench-dried inside climate chambers which were either set to a temperature of 25 °C and a vapour pressure deficit (VPD) of ∼1.9 kPa (for the control treatment) or to 45 °C and ∼6.7 kPa (for the heat treatment). One leaf per plant was fitted with an optical imaging set-up composed of a Raspberry Pi computer and a digital Raspberry Pi Camera Module V2 (Cavicam Co., Hobart, Tas., Australia; cavicam.com.au). Another cavicam was fitted in the closest available stem segment to monitor stem shrinkage over time. The cavicams also have a sensor that recorded the air temperature and humidity at 5-minute intervals, from which we calculated the ambient VPD. Once the cavicam system was set up, the sapling aboveground tissue was cut from the root, the cut was then sealed with white glue and images of both the target leaf and stem were captured every 5 minutes as the samples dried.

*Germany:* whole plants of the three study species were laid on benches within the climate chambers (York Industriekälte GmbH & Co. KG, Germany), which were set to either 20 °C (control treatment) or 35 °C (heat treatment) at 50% relative humidity, resulting in a VPD of

1.17 and 2.81 kPa, respectively. One leaf per plant was fitted with a custom-built optical imaging set-up composed of a Raspberry Pi computer and a digital Raspberry Pi Camera Module V2 (9132664, Raspberry Pi Ltd., Cambridge, UK) embedded in a custom-built 3D printed clamp (www.cavicams.com). Images were captured every 5 minutes as the samples dried.

### Image processing

All leaf image sequences from both Australia and German datasets were assessed using the well-established semi-automated image-processing pipeline for the OVT (www.opensourceov.org). Briefly, images were processed using FIJI software v2.16.0 (Schindelin et al., 2012) and the OSOV toolbox plugin (www.opensourceov.org). An image subtraction was performed, taking each image away from the one before to reveal the changes between the images (and therefore the embolism events). A series of steps was then undertaken to remove ‘noise’, i.e. the pixel signals from other changes that occur during drying, such as leaf movement and shrinkage, thereby isolating the embolism events. Embolism throughout the sequence was then expressed as a percentage of the total amount of pixels classified as part of embolism events observed as the sample dried, and plotted against water potential to construct vulnerability curves. All samples of *E. argophloia* and two samples from *Prunus avium* had to be removed from the dataset due to poor data quality resulting from incomplete and inconsistent detection of embolism events in the major veins.

### Water potential measurements

In both experiments, water potentials were measured with the pressure chamber method. Before excision, we bagged the leaves with aluminium foil for at least 30 min to ensure equilibration with the stem water potential (Rodriguez-Dominguez et al., 2022). We then measured water potentials using a Scholander pressure chamber (PMS Instruments, Corvallis, OR, USA). The first measurement was performed directly after placing the samples into the climate chambers. The subsequent measurements were done at different intervals in each branch of the experiment.

In the Australian branch, water potentials were initially obtained in 15–30 min intervals for the first few hours, which was increased to 1–3 h at more negative water potentials (no overnight water potential measurements were conducted). Typically, 6–15 (mean = 9) water potential measurements were obtained per treatment, except for *E. camaldulensis* in the heat treatment, where only 3–5 water potential measurements were obtained per sample, as this species dried down much faster. The stem diameter shrinkage and the corresponding leaf water potential measurements were then used to create a temporal series of water potential data for the entire experimental period using leaf optical dendrometry (Bourbia & Brodribb, 2023) assuming linear calibration relationships. The R^2^ of these leaf-specific calibrations varied from 0.71 to 0.99 with an average of 0.91 (Fig. S2).

In the German branch, water potentials were initially measured in intervals of 30–60 min, which were consecutively increased at more negative water potentials, resulting in median time differences between measurements of 1:33 h in the hot and 3:55 in the cold treatment. At more negative water potentials, we sampled two instead of one leaf to account for the higher variability between leaves in more dried-out plants. To interpolate water potentials for the vulnerability curves from Germany, we used shape constrained additive models based on `R` package `scam` v1.2-22 (Pya & Wood, 2015; Pya, 2026). Natural log-transformed (absolute) water potential was expressed as a function of time with a monotonically increasing P-spline, resulting in models with an R^2^ between 0.801 and >0.999 with an average of 0.967 (Fig. S3). Water potentials for all measurement times were then interpolated based on the back-transformed predictions from the additive models with a correction for retransformation bias (Smith, 1993).

### Data analysis

All data handling and statistical analyses were performed in `R` v4.5.2 (`R` Core Team, 2025) in the framework of the `tidyverse` (v2.0.0, Wickham et al., 2019).

VPD-corrected drying times (kPa h) for each sample were calculated as the cumulative sum of the product of the VPD for each time interval with its duration, thus weighting the elapsed time by the evaporative demand during that period (Blackman et al., 2019).

To analyse differences between water potentials associated with 12, 50 and 88% of cumulative embolism (*P*_12_, *P*_50_ and *P*_88_, respectively) between treatments (control vs. heat) in both datasets (Australian and German), we extracted these values from the raw observed vulnerability curves from both datasets, using the `approx()` function of base R to interpolate if required and averaging water potentials if more than one water potential value was observed for that particular value of cumulative embolism. In addition, we computed the expected embolism pressure (*P*_ex_) as the weighted average of the water potentials associated with embolism events weighted by the number of registered pixels in each event. This quantity can be considered the expected value of the distribution of embolism pressures described by the vulnerability curve, and has advantageous properties over *P*_12_, *P*_50_ and *P*_88_ because it quantifies a property of the entire curve instead of a single percentile (see Notes S1.1). Analogously, we extracted the VPD-corrected drying time until reaching 12, 50 and 88% percent cumulative embolism (*T*_12_, *T*_50_ and *T*_88_, respectively), and the expected value of the drying time distribution (*T*_ex_). Average vulnerability curves and cumulative embolism-drying time relationships were calculated by interpolating quantiles in regular intervals and averaging them. To quantify the uncertainty of average curves, we computed 95% parametric confidence intervals assuming approximate normality.

As *P*_12_, *P*_50_, *P*_88_ and *P*_ex_ are related properties determined from the same curves, they are highly correlated. To account for this non-independence, we analysed them with Bayesian multi-response mixed models using treatment, sampling location and their interaction as fixed effects, with residual correlations and treatment-specific random effects for species to account for the heterogeneity in the heat effects between species. We compared this model against a simplified model without site differences to estimate the overall treatment effect. Drying times were analysed with analogous Bayesian multi-response mixed models with *T*_12_, *T*_50_, *T*_88_ and *T*_ex_ as response variables. The models were fitted with the `R` package `brms` v.2.23.0 (Bürkner 2017; 2018) using moderately informative priors. Sampling was performed on 4 chains for 10000 iterations each, 5000 of which were discarded as warm-up. Convergence was evaluated based on the potential scale reduction factor 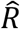 (Vehtari et al., 2021) and inspection of trace plots. Model adequacy was tested based on plots of posterior predictive checks (Gabry et al., 2019) and leave-one-out cross-validation based on `R` package `loo` v. 2.9.0 (Vehtari et al., 2017, 2025). Model comparisons were based on the expected log pointwise predictive density (ELPD) from leave-one-out cross-validation. We assumed parameter estimates to be credibly different from zero if their 95% credible intervals excluded zero. For details about model structure and prior specifications and derived quantities, see Notes S1.2 – S1.4.

## Results

### Observed vulnerability curves and effect of drying time

The average *P*_50_ of the curves differed between −2.91 ± 0.13 MPa for *Castanea sativa* and −6.43 ± 0.22 MPa for *Eucalyptus camaldulensis* (mean ± standard error), and was slightly less negative in the heat treatment compared to the control treatment for all studied species, though with a large variability within species (Fig. 3, Table 1). *P*_12_, *P*_88_ and *P*_ex_ showed analogous patterns (Fig. 3, Table 1).

**Table 1.**
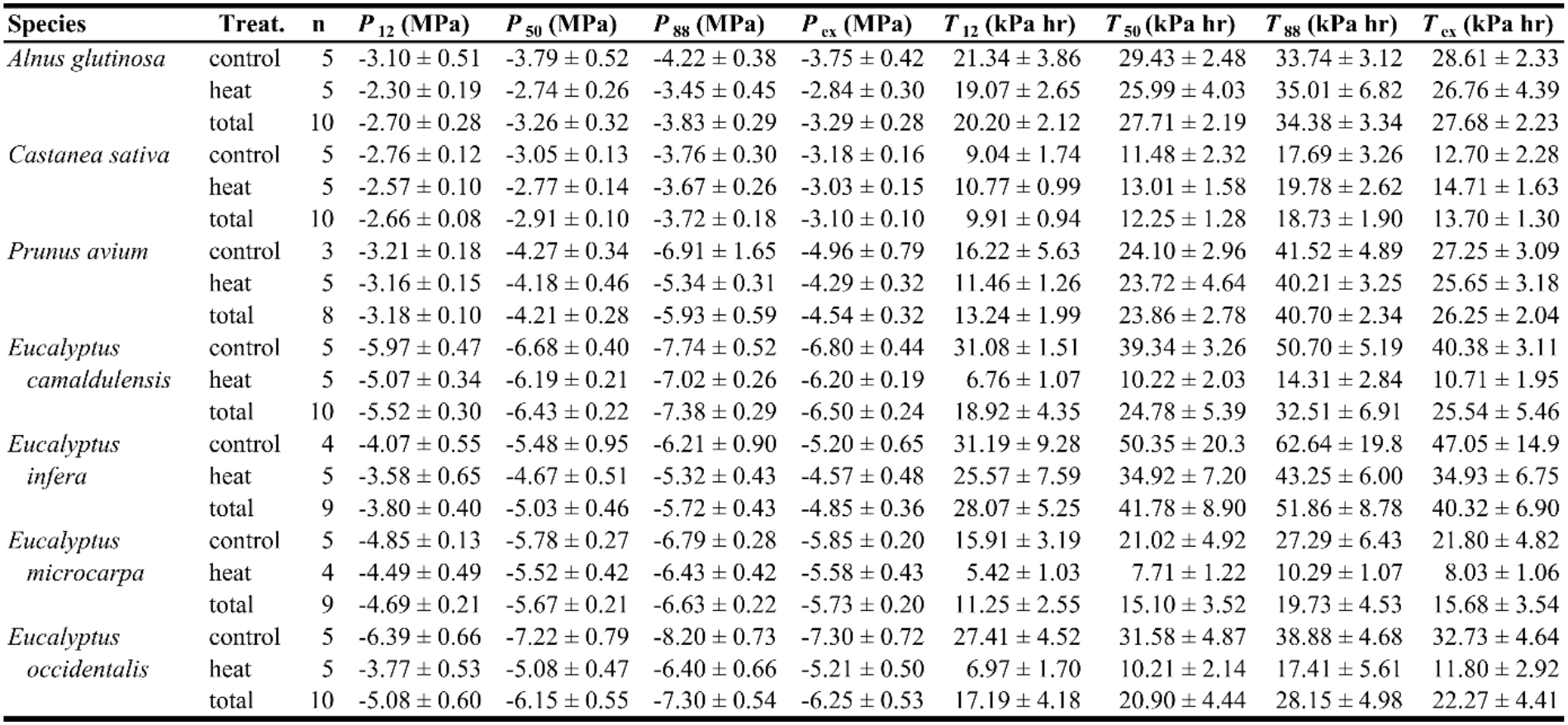
Observed water potential at 12, 50 and 88% percent cumulative embolism (*P*_12_, *P*_50_ and *P*_88_, respectively) and VPD-corrected drying time until reaching the water potential at 12, 50 and 88% percent cumulative embolism (*T*_12_, *T*_50_ and *T*_88_, respectively). Shown are mean ± standard error for the control (25 and 20 ºC for the Australian and German part of the experiment, respectively) and heat treatment (45 and 35 ºC for the Australian and German part of the experiment, respectively).

**Figure 3.**
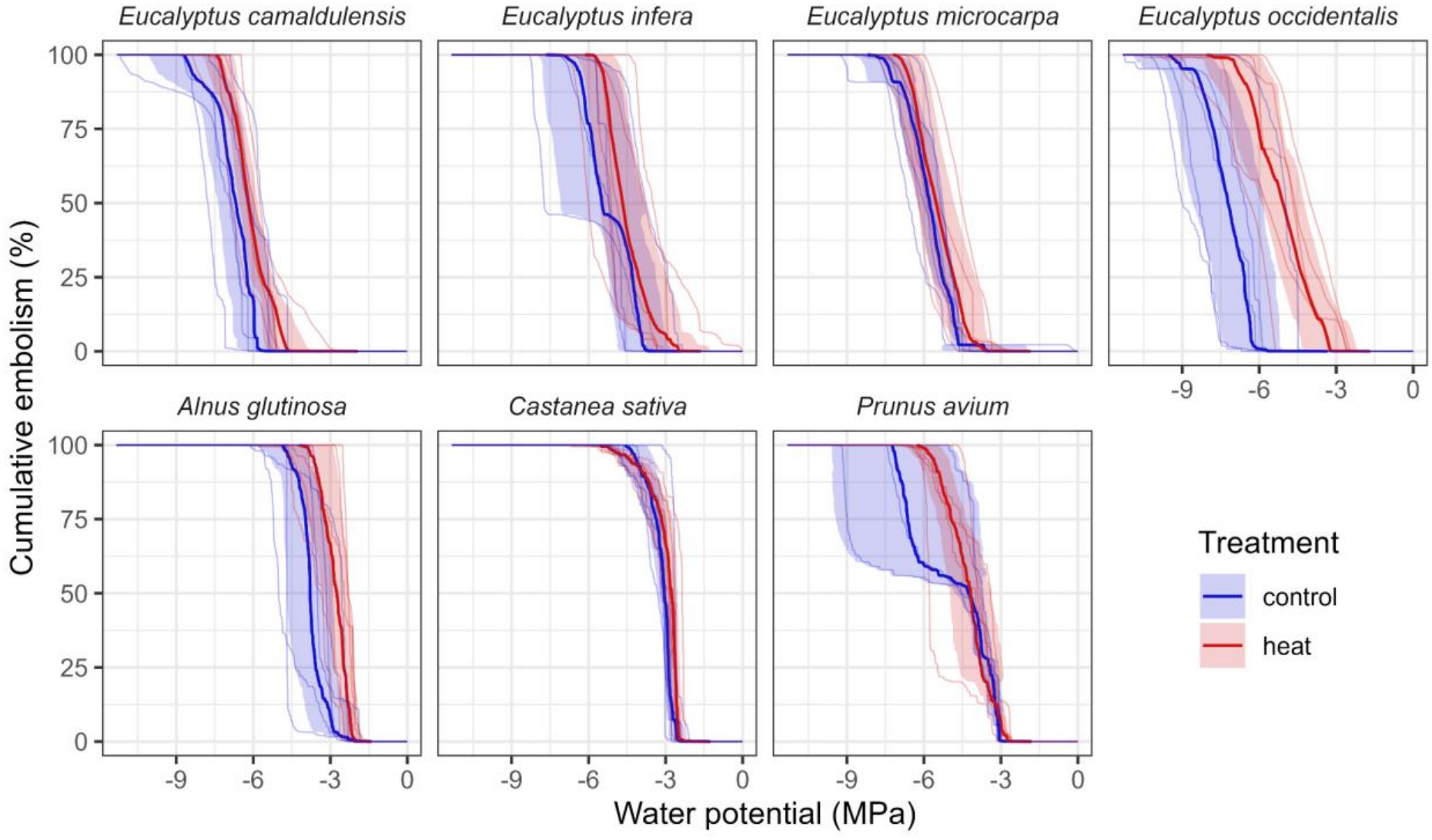
Vulnerability curves for the seven studied species in the control (blue) and heat treatment (red). Shown are the observed vulnerability curves obtained with the Optical Vulnerability Technique (faint lines) overlaid with the average curve (thick lines) with parametric 95% confidence intervals (shaded areas). Temperatures were set to 25 ºC (Australia) and 20 ºC (Germany) in the control and to 45 (Australia) and 35 ºC (Germany) in the heat treatment.

The average drying time until reaching the *P*_50_ differed greatly between species, ranging from 12.25 ± 1.28 kPa h for *Castanea sativa* to 51.86 ± 8.78 kPa h for *Eucalyptus infera* (Table 1). In contrast to the critical water potentials, for the drying times there was no consistent pattern in the treatment averages (Fig. 4), with consistently lower drying times at higher temperatures only observed in the samples from Australia (Table 1).

**Figure 4.**
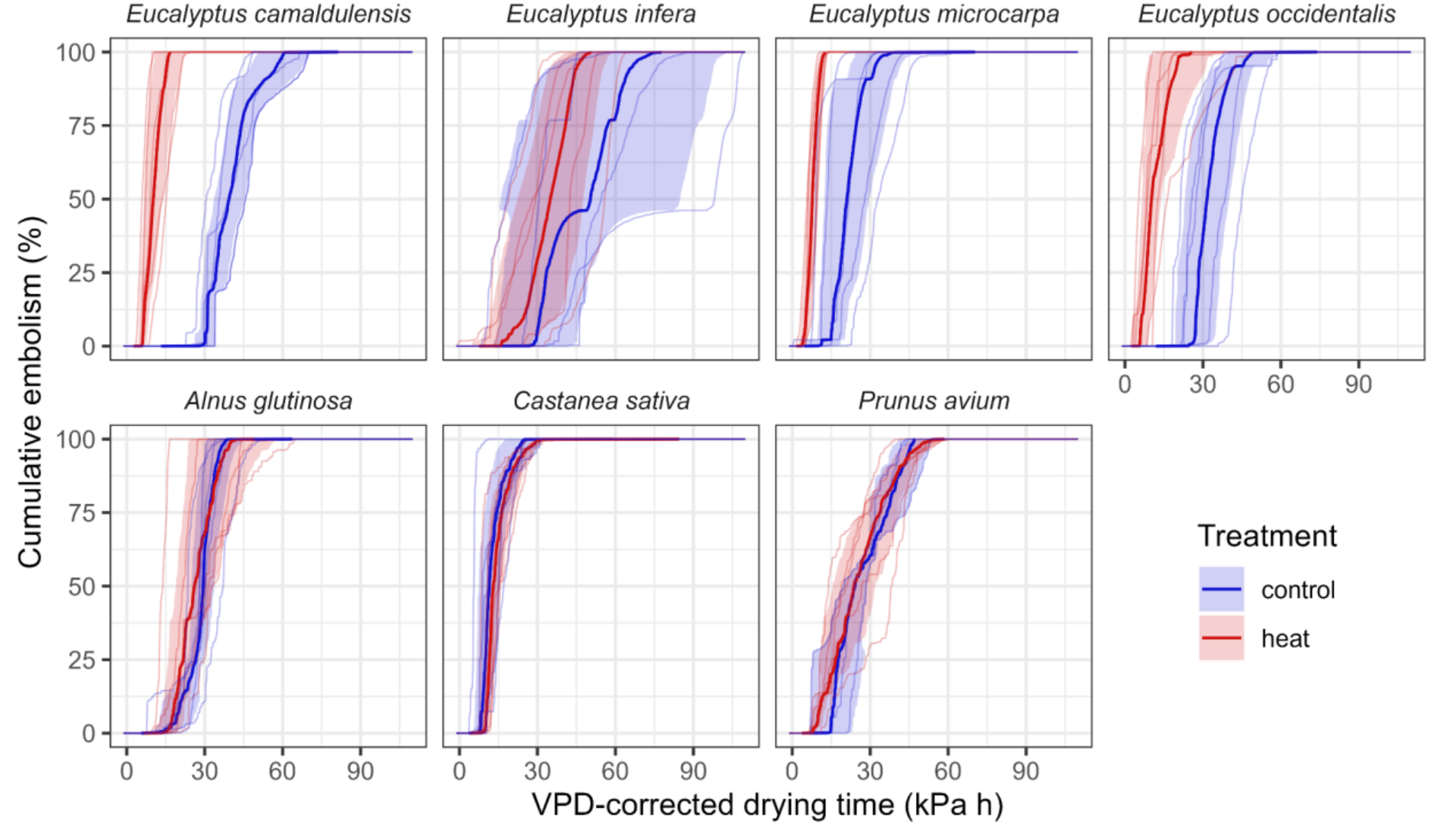
Change in cumulative embolism with VPD-corrected dehydration times for the seven studied species in the control (blue) and heat treatment (red). Shown are the observed values (faint lines) overlaid with the average curve (thick lines) with parametric 95% confidence bounds (shaded areas). Temperatures were set to 25 ºC (Australia) and 20 ºC (Germany) in the control and to 45 (Australia) and 35 ºC (Germany) in the heat treatment.

### The effect of heat on embolism thresholds

We found an effect of heat exposure on the water potentials at 12, 50 and 88% loss of conductivity and the expected embolism pressure, with the plants in the heat treatment reaching *P*_12_, *P*_50_, *P*_88_ and *P*_ex_ at credibly less negative water potentials than the control plants (Fig. 5, Fig. 6). On average, plants in the heat treatment had a *P*_12_, *P*_50_, *P*_88_ and *P*_ex_ that differed from the control treatment by +0.79 MPa, +0.73 MPa, +0.84 MPa and +0.75 MPa, respectively (Table S1).

**Figure 5.**
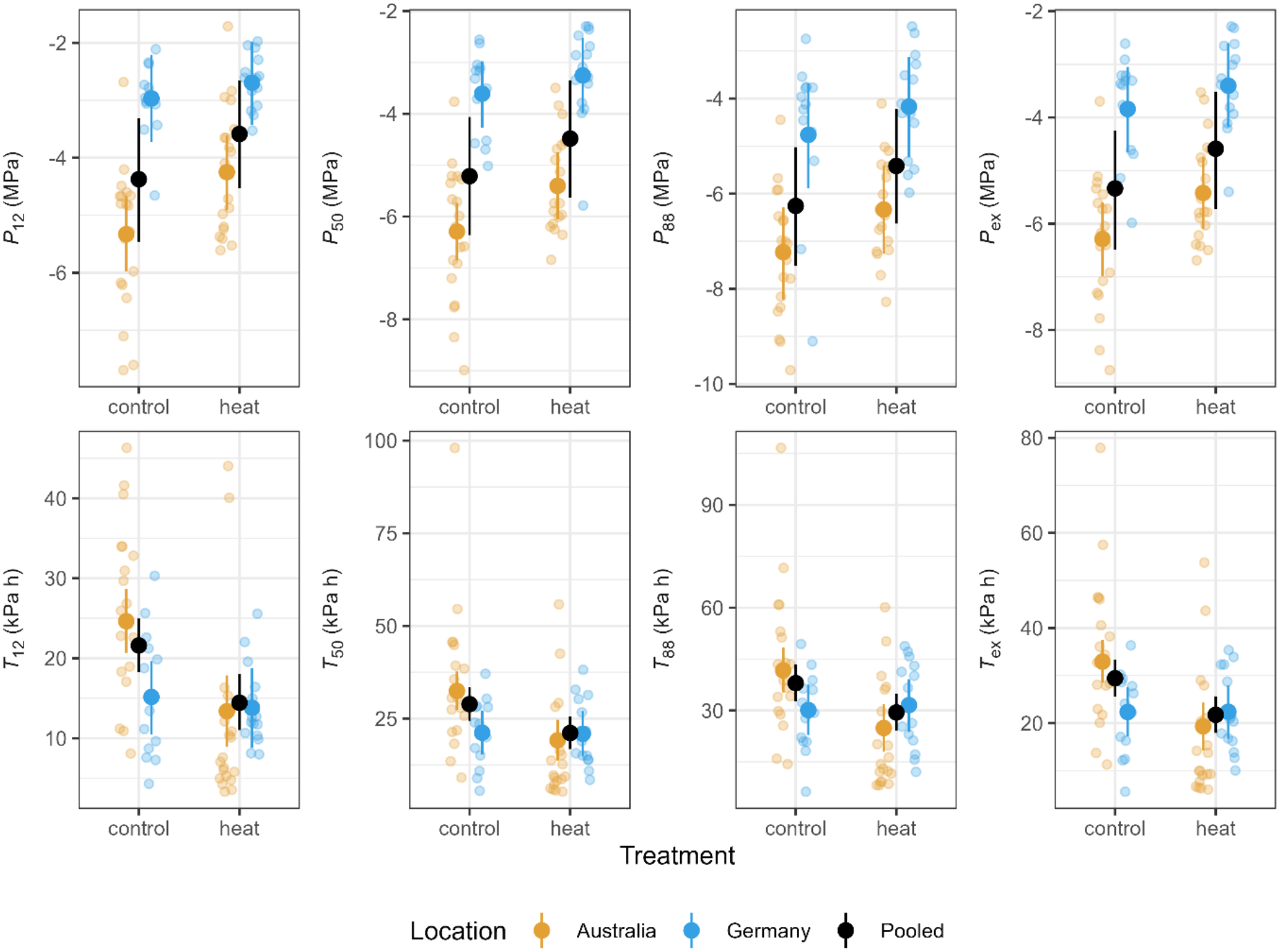
Water potential at 12, 50, 88% percent cumulative embolism and expected embolism pressure (*P*_12_, *P*_50_, *P*_88_, and *P*_ex_, respectively) and VPD-corrected drying time until reaching those water potentials (*T*_12_, *T*_50_, *T*_88_, and *T*_ex_) against treatments. Shown are the observed values coloured by location overlaid with the posterior means and 95% credible intervals for the marginal predictions of the models with pooled and site-wise treatment effects.

**Figure 6.**
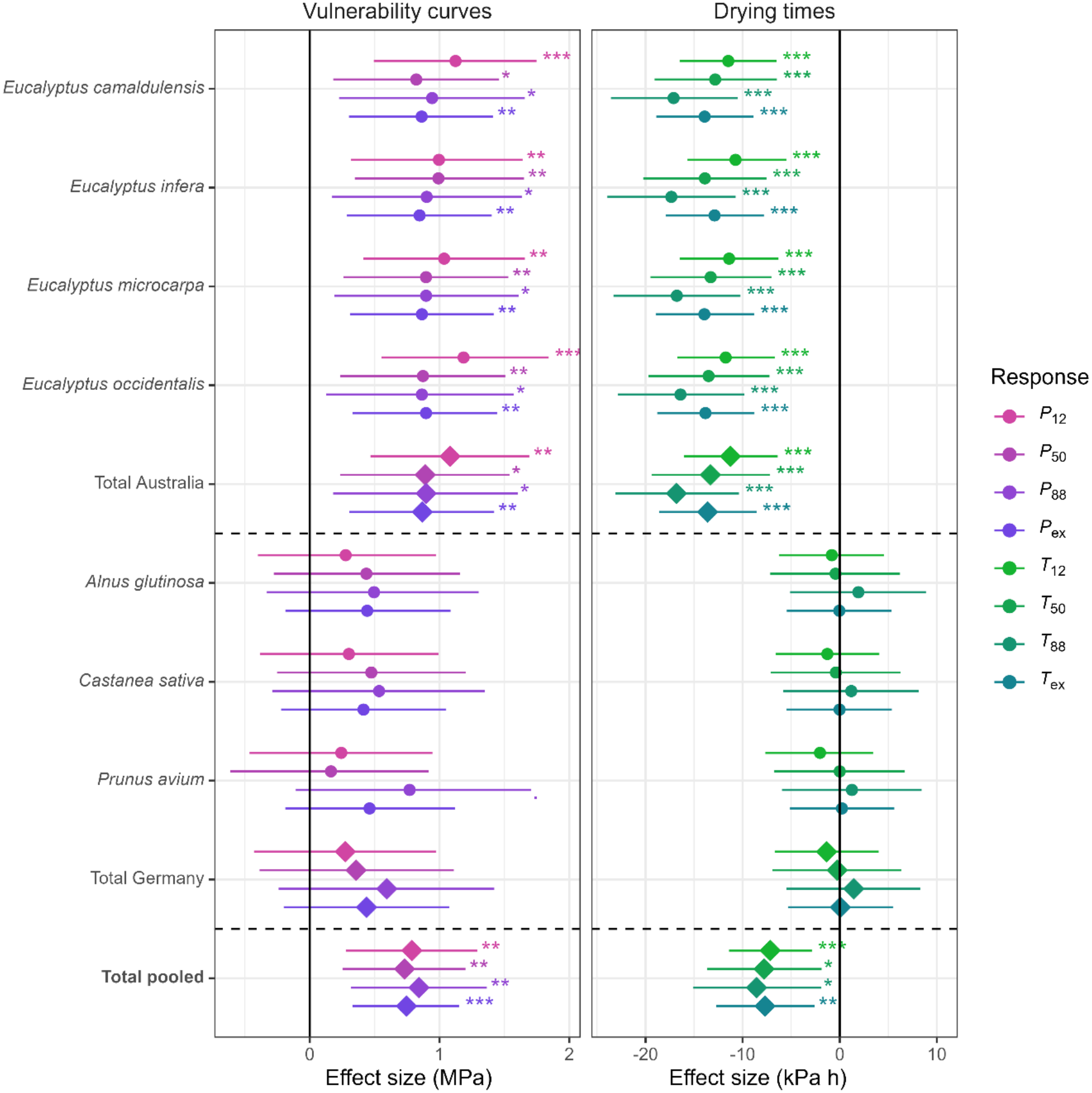
Effect sizes from models of vulnerability curve results (left) and drying times (right). Shown are the magnitude of the heat effects (posterior mean with 95% credible intervals) for the different species, as well as the estimates of the total effect for the two study locations, and pooled over both locations. Species-wise predictions are conditional effects of the site-specific model, total effects are based on the marginal effects of the site-specific (totals for Australia and Germany) and pooled model (pooled total). Symbols indicate credible differences on a 0.1 (.), 0.05 (*), 0.01 (**) and 0.001 level (***).

However, this effect was almost exclusively driven by the *Eucalyptus* species from the Australian branch of the study, while there was no conclusive evidence for a heat effect for the German plants (Fig. 6). The effect sizes for Australian species were even higher than in the pooled dataset, with increases in *P*_12_, *P*_50_, *P*_88_ and *P*_ex_ by +1.08 MPa, +0.89 MPa, +0.90 MPa and +0.87 MPa, respectively (Table S2). For the experiments in Germany, the heat treatment effect on critical water potentials neither differed credibly from zero nor from the effect size in Australia. In consequence, the pooled model had a slightly better fit to the data than the model with different means for Australia and Germany with a ΔELPD of 3.6, though with high standard error of 4.2, indicating that the two models were approximately equivalent.

### The effect of heat on drying time

The time until reaching 12, 50 and 88% loss of conductivity as well the expected value of the drying time distribution on average were all credibly lower for the plants in the heat treatment compared to the plants in the control treatment (Fig. 5, Fig. 6). On average, plants in the heat treatment reached their *T*_12_, *T*_50_, *T*_88_ and *T*_ex_ earlier by −7.2 kPa h for *T*_12_, by −7.8 kPa h for *T*_50_, −8.6 kPa h for *T*_88_ and by −7.7 kPa h for *T*_ex_ (Table S3).

As for the vulnerability curve parameters, the overall heat effect on the drying times resulted from the large effect sizes for the plants of the Australian branch of the study, while drying times nearly did not differ between treatments for the German plants (Fig. 6). The effect sizes for Australia were substantial, with *T*_12_, *T*_50_, *T*_88_ and *T*_ex_ reduced by −11.3 kPa h for *T*_12_, by −13.3 kPa h for *T*_50_, −16.8 kPa h for *T*_88_ and by −13.6 kPa h for *T*_ex_ (Table S4). Unlike the model for vulnerability curve parameters, in the model of drying times, the magnitude of the heat effect was credibly higher for the Australian species. Here, the grouped model had a better fit than the pooled model with a ΔELPD of 5.5 ± 6.3, although the large standard error again did not allow to rule out equivalence of the models.

## Discussion

### Heat exposure results in reduced embolism resistance

Our results provide evidence for a shift in vulnerability curves in a subset of species exposed to higher temperatures that supports our first hypothesis, with heat-exposed plants on average reaching critical levels of embolism at less negative water potentials. This is consistent with the proposed mechanism of temperature changes that shift the thermodynamic equilibrium in previously stable bubbles in the xylem sap to an unstable state that favours their expansion. However, there are other possible explanations for the observed decrease in embolism resistance. Based on the classical air-seeding concept, higher temperatures would be expected to reduce air-seeding pressures due to decreases in surface tension (Cochard, 2021). However, the observed effect size in our dataset is much larger than the +0.2% change per °C predicted by Cochard (2021). For the Australian species with an average *P*_50_ of −6.29 MPa, the increase in *P*_50_ of +0.89 MPa for a 20 °C increase in temperature was over 3.5 times higher than the 4% (+0.25 MPa) predicted based only on surface tension effects on air-seeding pressure. For the German species, which were subjected to a temperature difference of 15 °C, the mismatch in effect size was similar (+0.36 MPa instead of the expected 3% or +0.11 MPa), albeit here the credible intervals of the effect included zero. Accordingly, the effect of a temperature-driven decrease in surface tension on air-seeding pressures alone is not sufficient to explain the observed shift in vulnerability curves.

On the other hand, the temperature change alone (20 and 15 °C above ambient in the Australia and Germany, respectively) would increase *f*(*n*_*g*_), the effect of gas content on bubble stability (Eqn 10), by 22% percent in the Australian and 16% in the German experiment. This could push pre-existing nuclei in the sap, such as stable nanobubbles, into a state where no energy barrier exists for their expansion (Fig. 2), assuming that they were in equilibrium before heat exposure. Over a longer timescale, the reduced gas solubility at higher temperatures would promote gas diffusion into existing nuclei, further driving them into an unstable state. In xylem sap, nanobubbles are assumed to be coated with surfactant shells that promote their stability via dynamic surface tension (Schenk et al., 2015, 2017; Yang et al., 2020; Ingram et al., 2021). This means that bubble expansion leads to an increased surface tension due to the reduced density of surfactants on the larger surface, which then results in a new and larger stable radius (Schenk et al., 2017). This process is likely to provide a safety mechanism against the spontaneous bubble expansion that otherwise would be likely to happen after steep temperature increases due to the resulting changes in bubble stability. However, surfactant-coated bubbles are not exempt from the process described here, and radius-dependent surface tension only protects against bubble expansion until reaching values close to the surface tension of pure water (Yang et al., 2020). Extreme temperature changes may therefore also promote bubble nucleation in the presence of surfactants.

The changes in water viscosity related to the treatment are unlikely to have affected our results, as viscosity does not play a direct role in bubble nucleation (compare Eqn 1–18). While an increased hydraulic conductance resulting from lower viscosity (Cochard et al., 2000) to a certain degree may offset the loss of hydraulic conductance due to embolism, this does not play a role during the experimental dry-down of plants for the optical vulnerability technique because during the dry-down actual flows in the xylem are likely negligible, and because the method is not based on conductance measurements.

We observed a difference in the heat effect on vulnerability curves between the German and Australian branch of the experiment, with smaller effect sizes and no credible overall effect in Germany. A possible explanation is that the experimental supersaturation of xylem sap for the plants in Germany did not affect embolism formation, and that the larger temperature differential and higher end temperature in Australia had a larger impact. Since xylem sap is likely close to full gas saturation most of the time (Schenk et al., 2016), a low impact of artificial saturation is not surprising. However, we cannot exclude that the high temperatures of 45 °C in the Australian heat treatment may have caused leaf tissue damage resulting in an earlier failure of leaf veins that contributed to the observed heat effects. As the affected species come from regions in Australia that can be exposed to temperatures equal or even higher than 45 °C, and no leaf scorching was observed in our experiment, we are confident that our proposed mechanism at least partly explains the observed heat effects on the vulnerability curves, despite the varying results between the experiments.

### Heat exposure can accelerate drying time beyond the effect of VPD

The observed temperature-driven shift in vulnerability curves towards less negative water potentials was associated with a reduction in average VPD-corrected drying times.

Accordingly, the plants in the heat treatment would have reached critical thresholds of cumulative embolism earlier if VPD was held constant. This difference was only driven by the *Eucalyptus* species, with no evidence for a difference in the German part of the experiment. Therefore, our results only partially support our second hypothesis, and suggest that heat effects on plant dehydration time are context-dependent. Assuming a constant rate of water loss per kPa h between treatments, the observed change in vulnerability to embolism after heat exposure would result in reduced VPD-corrected drying times. However, the heat treatment reduced *T*_50_ by an average of −41% in the Australian and under −1% in German species, while *P*_50_ increased by +14% in the Australian and +9% in German species. The disproportionally larger heat effect on drying time in Australian species indicates that differences in water potential thresholds alone are not sufficient to explain the observed differences in the time to reach these thresholds. A potential explanation for the larger-than-expected reduction in drying times in Australian samples could be temperature-dependent increases in cuticular conductance (Cochard et al., 2021). This process could lead to higher residual water loss if exceeding the temperature associated with a conformation change of cuticular waxes, and has been demonstrated across a wide range of species (Schreiber, 2001; Riederer, 2006; Schuster et al., 2016). Another explanation could be stomatal reopening to permit evaporative cooling (Diao et al., 2024), which has been demonstrated to occur at higher temperatures in several species of *Eucalyptus* independent of water access (Marchin et al., 2023) and hence could have accelerated the dry-down of the Australian species. Meanwhile, for the plants from the German experiment, the lower maximum temperature of 35 °C makes it less likely that temperature thresholds were crossed that result in phase changes in the cuticle or trigger stomatal reopening. Rather, here the heat exposure led to earlier and more complete stomatal closure compared to the control treatment, resulting in no detectable difference in VPD-corrected drying time.

In recent years, a growing number of studies has assessed the factors that determine the timing to critical levels of dehydration (Blackman et al., 2019; Paligi et al., 2024, 2026; Waite et al. 2024, 2026). The optical vulnerability technique is a useful tool for studying drying times, as it results in high-resolution data of temporal changes in both water potential and the degree of embolism, and can be used across multiple plant compartments simultaneously (Harrison Day et al., 2024; Rodriguez-Dominguez et al., 2018). The differences in results between water potential thresholds and the times to reach them indicate that focusing on the temporal aspect of the dry-down instead of only reporting water potentials allows drawing a more complete picture of plant drought responses.

### Consequences for future studies of plant vulnerability to embolism

Our framework and experimental evidence indicate that critical water potential thresholds like *P*_12_, *P*_50_ and *P*_88_ may not be static traits, but instead may vary dynamically based on measurement conditions such as temperature and the gas content of the xylem sap. This has important repercussions for vulnerability curve measurements. Firstly, our findings highlight that for comparability of measurements obtained with the same method, measurement temperatures should be kept constant and should be reported together with the results. At least for the flow-centrifuge method, temperature-related changes in viscosity are known to be an issue affecting data quality (Wang et al., 2014; Burlett et al., 2022). However, we are not aware of studies that pointed out mechanisms how temperature could affect embolism formation itself with the exception of Cochard et al. (2021). Secondly, our findings indicate that the gas content of the liquid in the xylem can affect measurements. For that reason, methods where embolism formation happens in natural xylem sap, such as the optical vulnerability technique (Brodribb et al., 2016), classical bench dehydration (Sperry et al., 1988), micro-CT (Cochard et al., 2015), or the pneumatic method (Pereira et al., 2016) may come to systematically different results than methods that use a filtered and often degassed artificial measurement solution, such as the flow-centrifuge method (Cochard et al., 2005), or the air injection method (Cochard et al., 1992; Ennajeh et al., 2011).

An important consequence is that water potential thresholds obtained under laboratory conditions may not always reflect the behaviour of plants under natural conditions. During extreme heatwaves, plants may die *before* they actually reach the critical water potentials measured in the laboratory at room temperature. This can be relevant, for example, in modelling settings where embolism resistance measurements from global databases, like the Xylem Functional Trait Database (Choat et al., 2012) or TRY (Kattge et al., 2020), are used as static thresholds for modelling drought-induced plant mortality under rising temperatures. If the mechanism outlined here—in which embolism develops at lower water potentials than previously expected—is confirmed in further studies, this would have significant implications for assessing the drought tolerance of tree species in a changing climate.

While our experimental evidence is in line with the idea that rapid temperature changes can reduce the stability of bubbles in xylem sap and thus lead to increased xylem vulnerability, we are aware of the pilot-study character of our work. We acknowledge that our study combines results from two small experiments with limited sample sizes and differing in design, and that more rigorous testing of the mechanism in larger studies is necessary. Future studies should test for temperature effects on xylem vulnerability curves using a larger number of species and larger sample sizes, using consistent heat treatments and a fully-factorial design for evaluating the effect of experimental supersaturation. As direct evidence of embolism formation in the xylem is nearly impossible to obtain, this should be accompanied with mechanistic modelling of bubble nucleation processes in the xylem that take into account the influence of gas content.

## Conclusions

The theoretical framework presented here highlights the importance of dissolved gas for bubble nucleation processes in xylem sap, describes the conditions under which gas-filled nuclei can remain stable, and shows how sudden temperature changes can result in the spontaneous expansion of formerly stable nuclei. The effect of gas content on bubble stability adds an important new piece to the puzzle of the mechanisms of xylem embolism formation. Our experimental evidence suggests that in line with the proposed mechanism, heat exposure during drought can result in reduced embolism resistance and a faster dry-down even after correcting for differences in VPD, at least in a subset of species and at large enough temperature differences. Our framework may thus help explaining the excessive tree mortality observed following hotter-drought events.

## Acknowledgements

This study was supported by the ARC-DECRA 2026 Fellowship (DE260100224), the Bert and Aileen Kollosche Bequest research grant 2025 from the Australian Plant Society of South Australia, and by the Thomas Davies Research Grant for Marine, Soil and Plant Biology 2026 from the Australian Academy of Sciences awarded by ISM. KMJ was funded by the European Union on a Marie Skłodowska-Curie Actions (MSCA) Postdoctoral fellowship, 101107177— IVERdrought. Views and opinions expressed are, however, those of the author(s) only and do not necessarily reflect those of the European Union or the MSCA. Neither the European Union nor the granting authority can be held responsible for them. KMJ acknowledges the travel grant provided by the University of Gottingen’s faculty of forest sciences and forest ecology which facilitated collaboration including the experimental work in Germany. SSP and CA acknowledge the funding support which enabled the establishment of the pot experiment in Göttingen, Germany through grant number FKZ 67KIA4059E under the KI-recover project. AR was supported by the DAAD fellowship to visit the department of Plant Ecology and Ecosystems Research and Silviculture and Forest Ecology of Temperate Zones, University of Göttingen, Germany. We gratefully acknowledge Regina Helbig, Heinz Coners and Bharath Kumar Haralalli Nagarajappa for technical support in establishing and maintaining the potted saplings at the Experimental Botanical Garden in Göttingen, Germany. RML and KP wish to thank BS (R.I.P.) & G for moral support.

## Author contributions

RML and ISM developed the original idea of the project, which was then considerably modified in collaboration with KMJ and SSP. RML developed the theoretical framework. ISM planned the Australian experiment and collected the Australian dataset together with NA, JDKD, and WRW.SSP, KMJ and RML planned the German experiment and collected the German dataset together with KP, AR and FRT. RML wrote the original draft with help from SSP, KMJ and ISM, which was then revised and approved by all other authors.

## Data availability statement

Upon publication, the dataset and R code underlying the analysis in this paper will be made available as a public repository on zenodo.org.

## Boxes

### Box 1

**Temperature influence on the behaviour of xylem sap**

Several physical properties describing the gas-water interface in the xylem are temperature dependent. For embolism formation, the most relevant processes after an increase in temperature are an increase in gas solubility and a decrease in surface tension. These changes are accompanied by a decrease in water viscosity.

*Gas solubility*

The solubility of gas in water can be described by Henry’s law:

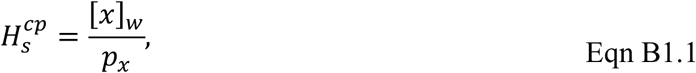

where 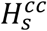 is the Henry’s law constant, [*x*]_*w*_ is the equilibrium concentration of the gas *x* in water (mol L^−1^), and *p*_*x*_ is its partial pressure of *x* in air.

The temperature dependence of equilibrium constants like 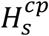 is described by the Van’t Hoff equation:

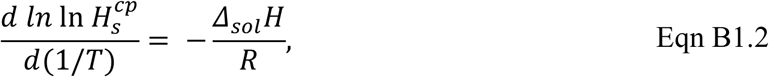

Where *T* is the temperature in Kelvin and Δ_*sol*_*H* is the enthalpy of dissolution (J mol^−1^). Integrating this equation and expressing 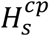 based on a reference value *H°* at the reference temperature *T°* = 298.15 K results in the following expression describing an exponential decrease in solubility with temperature with a rate determined by the enthalpy of dissolution:

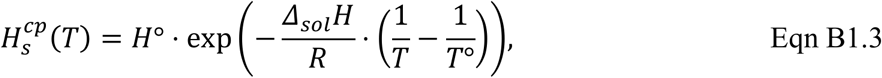

This equation is accurate only for a limited range of temperatures over which the enthalpy of dissolution is approximately constant with temperature. Values of *H°* and Δ_*sol*_*H*/*R* for a large range of liquids can be found in Sander et al. (2023).

*Surface tension*

The drop in gas solubility with increasing temperatures is accompanied by a reduction in surface tension. There are a large number of empirical equations to approximate the temperature dependence of surface tension. We present here the model by Vargaftik et al. (1983) in the revision by IAPWS (2014):

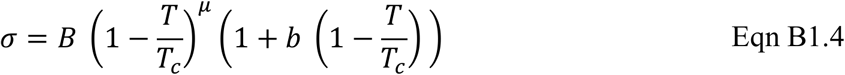

where *σ* is the surface tension (J m^−2^) of pure water, *T* is the absolute temperature in Kelvin, *T*_*c*_ is the critical temperature of water (647.096 K), and *B, μ* and *b* are constants (*B* = 0.2358 N m^−1^, *μ* = 1.256, *b* = 1.256). In the presence of surfactants, *σ* will be reduced, but show a comparable temperature dependence.

*Viscosity*

Increasing temperatures result in a decline in viscosity. A large number of empirical equations has been proposed for the relationship between the viscosity of water and temperature, one of which can be found in Kestin et al. (1978):

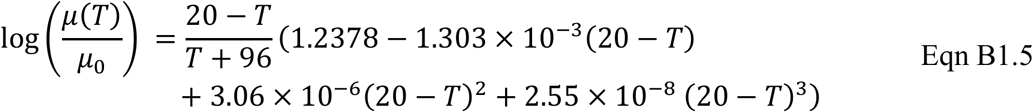

where *μ*(*T*) is the viscosity of the liquid (Pa s) at temperature *T* (°C), *μ*_0_ is the viscosity at a reference temperature of 20 °C (1.002 Pa s), which achieves an accuracy of ± 0.2% over the temperature range from −8 ≤ *T* ≤ 150 °C.

## Supporting information

### Notes S1. Supplementary methods

In this document, we first provide a derivation of the expected embolism pressure *P*_ex_ as a metric for comparing vulnerability curves (S.1.1), and then describe the structure of the multivariate model of different water potential thresholds (*P*_12_, *P*_50_, *P*_88_ and *P*_ex_) in dependence of heat treatment and study location (S1.2 – S1.4). The models for VPD-corrected drying time (*T*_12_, *T*_50_, *T*_88_ and *T*_ex_) followed an analogous structure and are hence not described separately.

#### S1.1 Expected embolism pressure P_ex_

To describe differences in embolism resistance, it is customary to compute point estimates of water potential thresholds such as the *P*_12_, *P*_50_ or *P*_88_, and then compare how they differ between curves. However, the thresholds of 12, 50 and 88% cumulative embolism are largely arbitrary, and if the curves are rather asymmetric, as is usually the case for the optical vulnerability technique, slight differences in the percentage of cumulative embolism can result in large differences in water potential. It would therefore be advantageous to have a metric to compare the location of curves across their entire range.

Vulnerability curves describe the change of cumulative embolism with decreasing water potential, and can thus intuitively be interpreted as cumulative distribution functions of the underlying distribution of embolism pressures. In the following, assume two arbitrary vulnerability curves *F*_1_(Ψ) and *F*_2_(Ψ) that both map water potentials Ψ onto the observed fraction of cumulative embolism *p*. A reasonable metric of distance over the entire curves is the expected value E[*d*] of the water potential distance *d* between the two curves over their entire range. Using the definition of the expected value of a continuous function *f*(*x*) over the interval [*a, b*] as 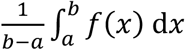 can be calculated as the integral over the difference of their quantile functions 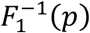 and 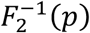:

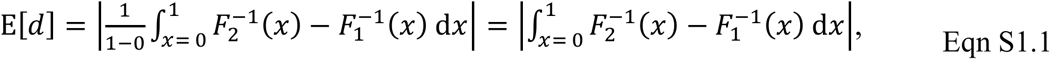

The integral in Eqn S1.1 can be split into two components, resulting in:

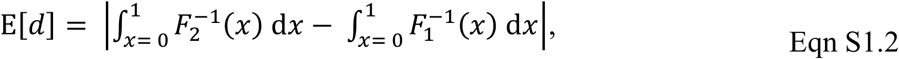

It can be shown that the integral over a quantile function is equal to the expected value of the corresponding distribution. Accordingly, assuming that *X*_1_ and *X*_2_ are random variables following the cumulative distribution functions *F*_1_(Ψ) and *F*_2_(Ψ), the expected water potential distance between curves reduces to:

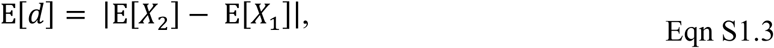

This means that the expected value of the water potential difference between two vulnerability curves is simply the difference between the expected values of the embolism pressure distribution described by the two curves. It follows that comparing differences in E[*X*] is analogous to comparing the expected water potential difference E[*d*] calculated over the entire range of the vulnerability curves. In this work, the quantity E[*X*] is denoted as *P*_*ex*_.

For symmetrical models of vulnerability curves such as the commonly used exponential-sigmoid model of Pammenter & Vander Willigen (1998), *P*_*ex*_ and *P*_50_ are identical, so that comparisons in *P*_50_ are consistent with comparing water potential differences along the entire curves. For the comparison of metrics from curves obtained with different statistical models, or for the highly asymmetric curves obtained by the optical method, this is not necessarily the case. Here, comparisons of *P*_*ex*_ instead of conventional thresholds like *P*_12_, *P*_50_ or *P*_88_ have the vast advantage of being equivalent to a comparison over the entire curve without actually having to evaluate the integral in Eqn S1.1.

For the optical vulnerability technique, *P*_*ex*_ can easily be computed as the weighted average of the water potentials Ψ for embolism event *i* weighted by the number of pixels *px*_*i*_ that were classified as embolized during this embolism event:

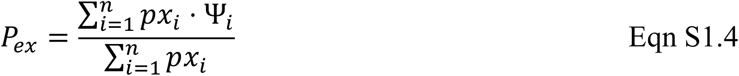

#### S1.2 Model structure

In the following, we will use the subscript *x* to mark statements that are valid analogously for all water potential thresholds considered in the model, e.g. *P*_*x*_ to signify all water potential thresholds (*P*_12_, *P*_50_, *P*_88_ and *P*_ex_). In our model, we described the joint distribution of the water potential thresholds *P*_*x*[*i*]_ for observation *i* with a multivariate normal distribution with mean *μ*_*x*[*i*]_ and 4 × 4 covariance matrix Σ_*res*_:

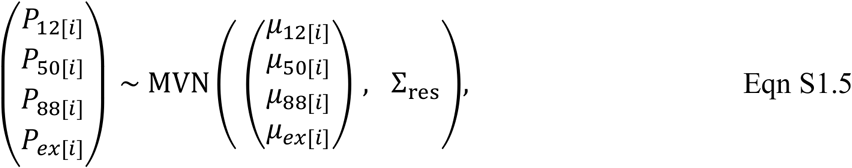

The mean *μ*_*x*[*i*]_ for each water potential threshold for species *j* and location *k* (Australia or Germany) was expressed as follows:

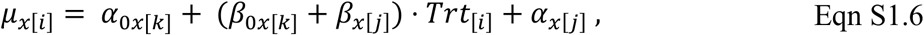

where *α*_0*x*[*k*]_ is the location-specific intercept, *Trt*_[*i*]_ is the heat treatment (control or heat) for observation *i, β*_0*x*[*k*]_ is the location-specific heat effect, and *α*_*x*[*j*]_ and *β*_*x*[*j*]_ are species-specific random intercepts and slopes, respectively.

We additionally fitted a simplified version of the model without location-specific differences:

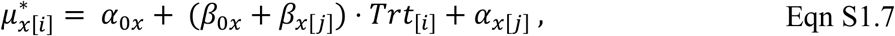

The random intercepts and slopes were described by a multivariate normal distribution centred around zero:

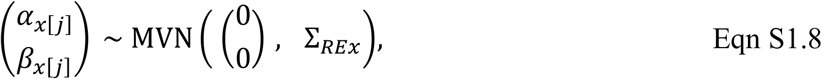

where Σ_*REx*_ is a 2 × 2 covariance matrix describing the joint distribution of the random effects, containing the random effects standard deviations *τ*_*αx*_ and *τ*_*βx*_ and the random effects correlation matrix Ω_RE_:

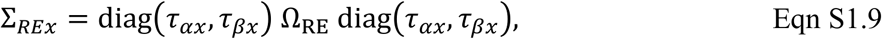

Analogously, the covariance matrix Σ_*res*_ of the residuals consists of the residual standard deviations *σ*_*x*_ and the residual correlation matrix Ω_res_:

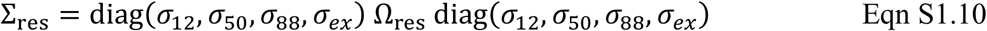

#### S1.3 Choice of priors

The location intercepts *α*_0*x*[*k*]_ (i.e. the average of the water potential threshold *P*_*x*_ in the control treatment at the corresponding location) were loosely centered around the observed average 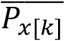 in the control treatment with a normal prior with a standard deviation of 2:

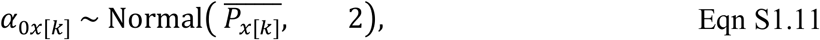

which puts 95% of the prior probability mass on values of approximately ± 4 MPa around the average in the control treatment. For the analogous models of drying time, due to the different scale of the response, the normal prior on the location intercepts was centered around the observed average 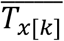 in the control treatment with a standard deviation of 10, which puts 95% prior probability mass on values of approximately ± 20 kPa h around the average in the control treatment. For the simplified version of the model without location differences, we used the pooled average 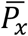 of the control treatment instead of 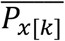.

The location-specific heat effects *β*_0*x*[*k*]_ were assigned normal priors with mean 0 and a standard deviation of 2:

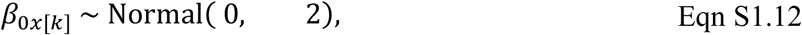

which centers the expected heat effects around zero, putting 95% prior probability mass for the difference between treatments on values of approximately ± 2 MPa. For the analogous models of drying time, the normal prior on the heat effects was centered around 0 with a standard deviation of 10, which puts 95% prior probability mass on values of approximately ± 20 kPa h.

The standard deviations of the random effects *τ*_*αx*_ and *τ*_*βx*_ were assigned gamma priors with shape 2 and rate 1 and 4, respectively:

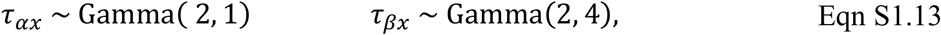

Gamma(2, ·) priors are boundary avoiding priors (cf. Gelman et al., p316) that discourage very low values for the random effect standard deviations. The priors put 95% probability mass on values between 0.24 and 5.57 with an expected value of 2.00 for *τ*_*αx*_ and on values between 0.06 and 1.39 MPa with an expected value of 0.50 MPa for *τ*_*βx*_, reflecting a higher plausible variability between species than between treatments. For the analogous models of drying time, we used a Gamma(2, 0.25) prior for *τ*_*αx*_ and a Gamma(2, 1) prior for *τ*_*βx*_, which have expected values of 8.0 and 2.0 kPa h and put 95% of the probability mass on values from 1.0 – 22.3 kPa h and from 0.2 – 5.6 kPa h, respectively. The use of boundary-avoiding priors for *τ*_*αx*_ and *τ*_*βx*_ was motivated by the low number of seven species in the dataset, which can affect the stability of the estimates of random effect standard deviations.

The residual standard deviations *σ*_*x*_ were assigned half-normal priors with a mean of 0 and a standard deviation of 2:

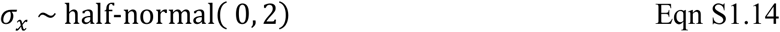

This prior puts 95% probability mass on values of *σ*_*x*_ below 3.3 MPa. For the analogous model of drying time, *σ*_*x*_ were assigned half-normal priors with a mean of 0 and a standard deviation of 10, which puts puts 95% probability mass on values of *σ*_*x*_ below 16.4 kPa h.

Finally, the correlation matrices Ω_RE_ and Ω_res_ were assigned LKJ(1) priors (Lewandowski et al., 2009), which are flat priors on correlation matrices that do not favour any specific values of correlations:

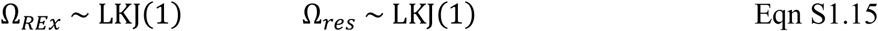

#### S1.4 Model fitting, validation and inference

The models were fitted in `R` v4.5.2 (`R` Core Team, 2025) using `R` package `brms` v.2.23.0 (Bürkner 2017; 2018). For all models, we used 4 parallel chains with 10000 iterations each. The first 5000 iteration were discarded as warm-up, resulting in a total of 20000 posterior draws for each model parameter. For all models, we set the target acceptance rate to 0.99 and used a maximum tree depth of 12.

Convergence was evaluated based on the potential scale reduction factor R (Vehtari et al., 2021) and inspection of trace plots. Model adequacy was tested based on plots of posterior predictive checks (Gabry et al., 2019) and leave-one-out cross-validation based on R package loo v. 2.9.0 (Vehtari et al., 2017, 2025), setting `re_loo = TRUE` to guarantee reliable estimates of the expected log pointwise predictive density (ELPD), since a low number of observations resulted in Pareto *k* values of above 0.7. Model comparisons were based on the ELPD from leave-one-out cross-validation.

As a Bayesian equivalent to the frequentist p-value, we computed the probability of direction *pd* for all fixed effects. The probability of direction *pd* is defined as the proportion of the posterior distribution that has the same sign as its median (Makowski et al., 2019).

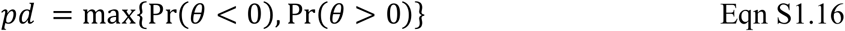

Based on *pd*, we then calculated a two-sided Bayesian pseudo-p-value *P*_*Bayes*_ as follows:

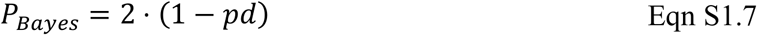

In a model with flat priors, *P*_*Bayes*_ is identical to the two-sided frequentist p-value (Makowski et al., 2019).

## Supplementary figures

**Figure S1.**
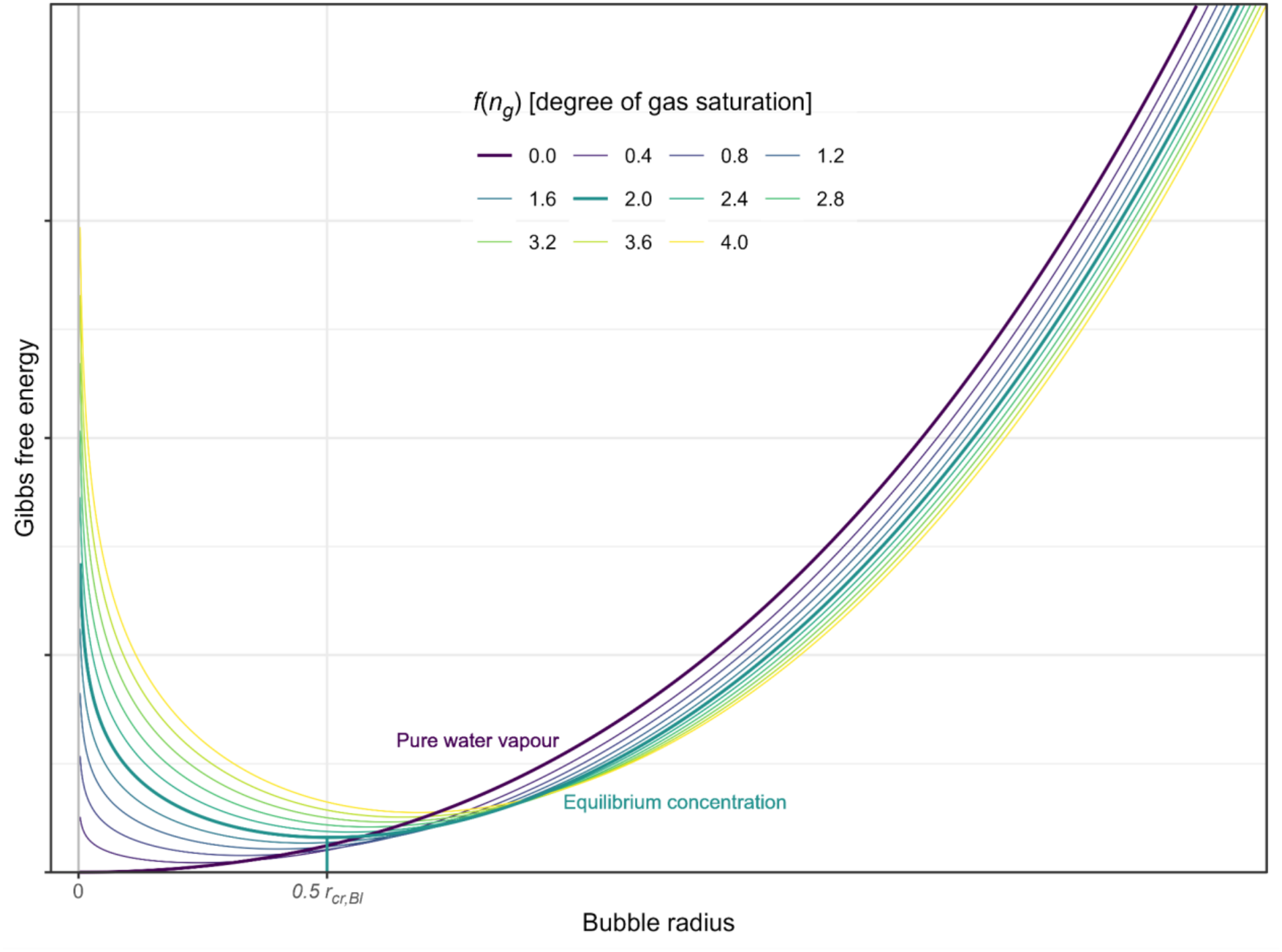
Change in Gibbs free energy with bubble radius for different gas contents *n*_*g*_ under the desinent regime. Shown are the (relative) values of 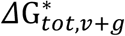following Eqn 7 coloured by *f*(*n*_*g*_) (an indicator of the degree of gas saturation of the gas in the bubble as a function of gas content, ranging from 0 for pure water vapour over 2 for equilibrium saturation to > 2 for supersaturated conditions; see Eqn 10). For pure water vapor, the equilibrium radius is zero. For all other gas contents, there exists an equilibrium radius (i.e., a local minimum), which is exactly half of Blake’s radius *r*_*cr,Bl*_ (Eqn 6) under equilibrium conditions (*f*(*n*_*g*_) = 2).

**Figure S2.**
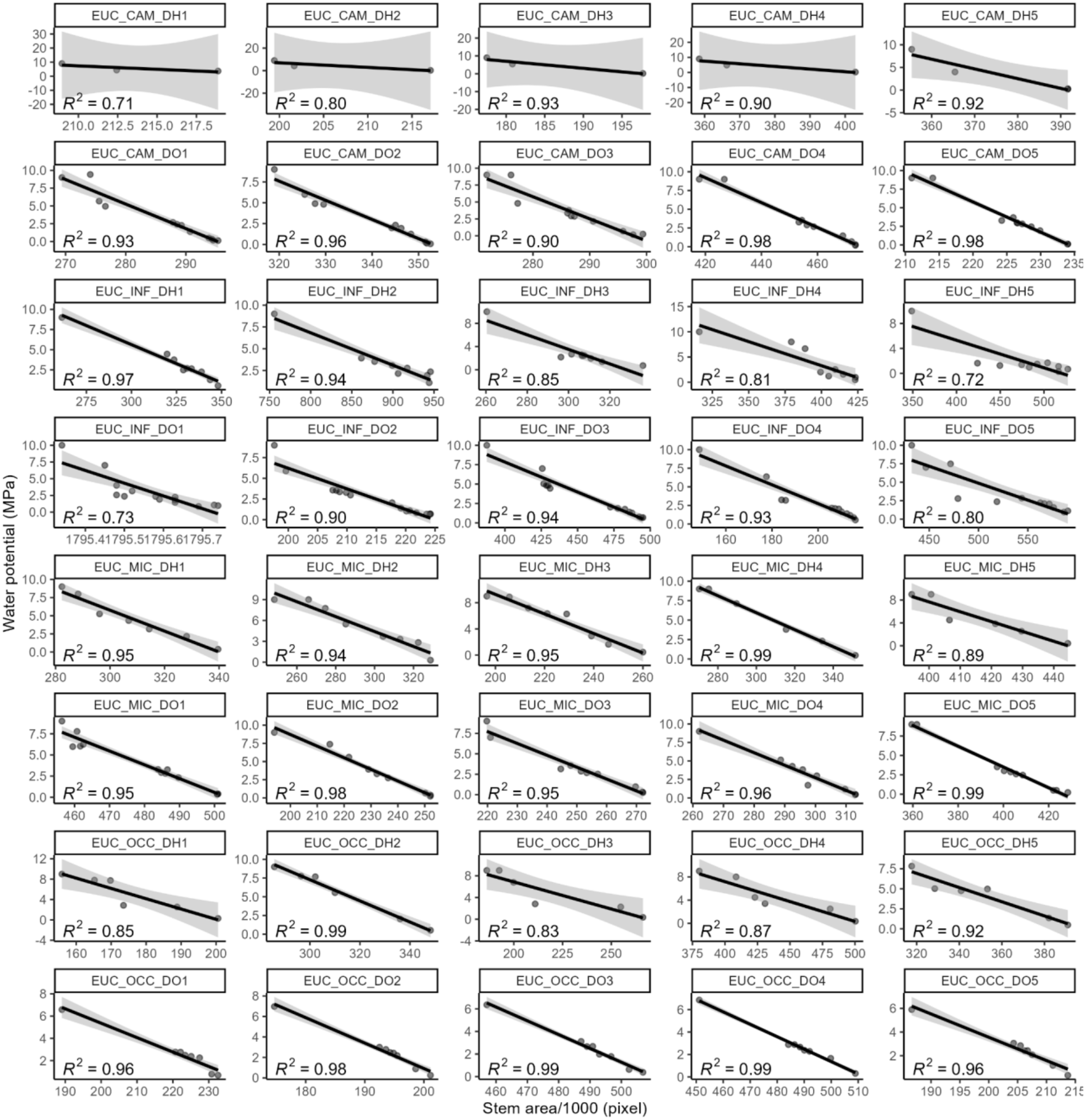
Calibration of water potentials for the Australian samples via leaf dendrometry for the control (DO1 – DO5) and heat (DH1 – DH5) treatments and four *Eucalyptus* species: *E. camaldulensis* – EUC_CAM, *E. infera* – EUC_INF, *E. microcarpa* – EUC_MIC, *E. occidentalis* – EUC_OCC. The models explained on average 91% of the variance in water potential, ranging from 71 to 99%.

**Figure S2.**
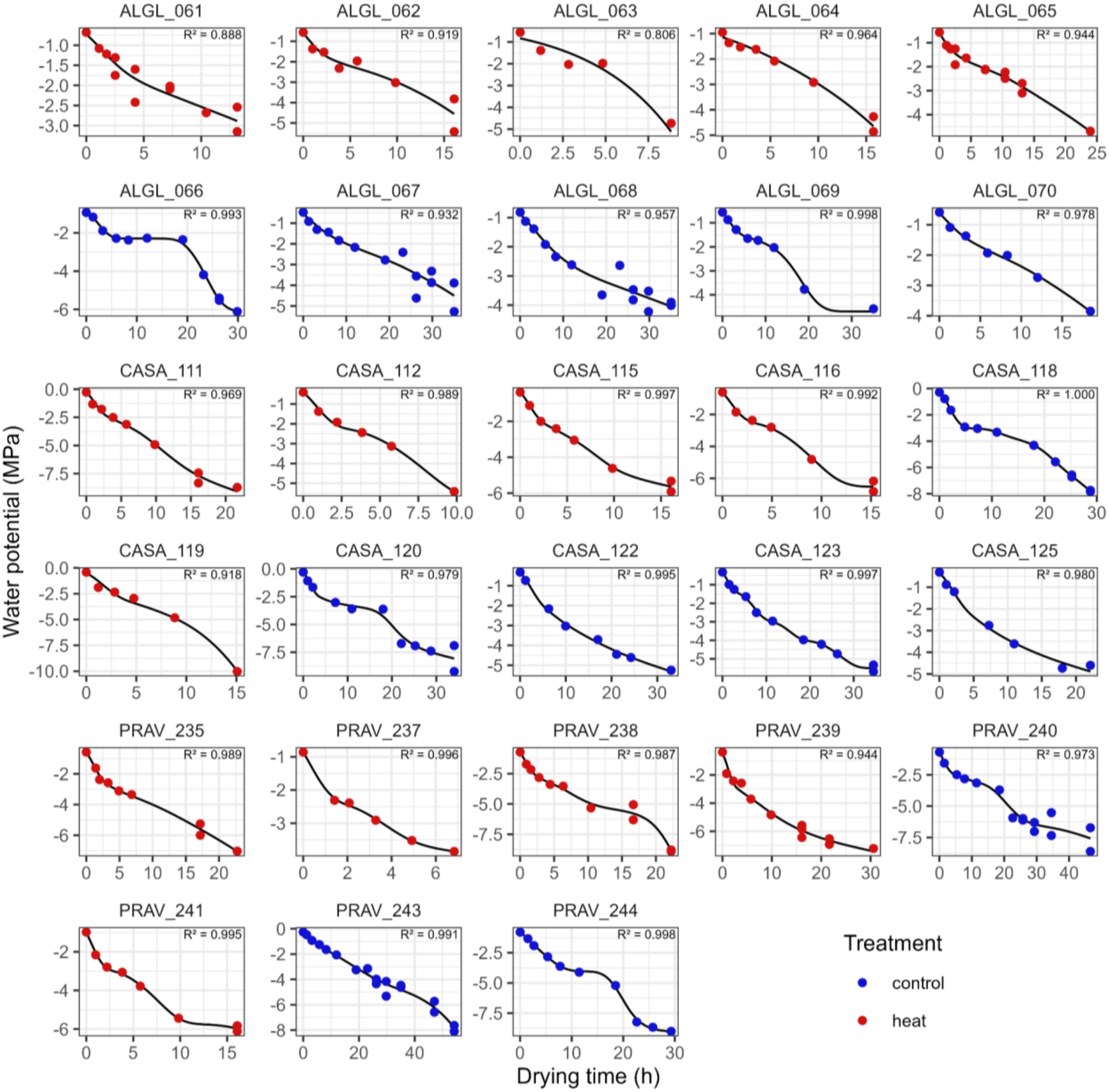
Relationship between (uncorrected) drying time and observed water potentials for the samples from Germany. Shown are the observed water potentials (points, coloured by treatment) and the predictions from the shape-constrained additive models (lines). Panel titles are sample IDs (*Alnus glutinosa* – ALGL, *Castanea sativa* – CASA, *Prunus avium* – PRAV). The models explained on average 96.7% of the variance in water potential, ranging from 80.6 to > 99.9%.

## Supplementary tables

**Table S1.** Results of the multi-response mixed model of *P*_12_, *P*_50_, *P*_88_, and *P*_ex_ pooled over both locations. See Notes S1.2 for model structure. Shown are the posterior means of the parameter estimates with their MCMC standard error, 95 % credible interval, potential scale reduction factor 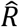, bulk and tail effective sample sizes and a Bayesian pseudo-p-value (only shown for fixed effects, see Notes S1.4). Symbols indicate parameters credibly different from zero on the 0.1 (.), 0.05 (*), 0.01 (**) and 0.001 level (***).

|  |  |  |  | **95 % CI** |  |  | **Eff. Sample size** |  |  |  |
|---|---|---|---|---|---|---|---|---|---|---|
| **Response** | **Parameter** | **Estimate** | **Est.Err.** | **lower** | **upper** | **R̂** | **Bulk** | **Tail** | ***P***<sub>**Bayes**</sub> |  |
| *P*<sub>12</sub> | *α*<sub>0 12</sub>: Intercept | -4.373 | 0.534 | -5.462 | -3.313 | 1.0007 | 6654.9 | 8830.8 | < 0.001 | *** |
|  | *β*<sub>0 12</sub>: Heat effect | 0.787 | 0.256 | 0.280 | 1.291 | 1.0002 | 10711.4 | 12472.0 | 0.004 | ** |
|  | *τ*<sub>*α* 12</sub>: RE SD Intercept | 1.335 | 0.457 | 0.697 | 2.444 | 1.0003 | 7840.0 | 10320.7 |  |  |
|  | *τ*<sub>*β* 12</sub>: RE SD Heat effect | 0.339 | 0.196 | 0.062 | 0.818 | 1.0004 | 9506.0 | 10745.7 |  |  |
|  | *ρ* (*τ*<sub>*α* 12</sub>, *τ*<sub>*β* 12</sub>): RE correlation | -0.634 | 0.403 | -0.995 | 0.521 | 1.0000 | 13617.8 | 13985.1 |  |  |
|  | *σ*<sub>12</sub>: Residual SD | 0.868 | 0.085 | 0.723 | 1.057 | 1.0005 | 9774.2 | 9726.6 |  |  |
| *P*<sub>50</sub> | *α*<sub>0 50</sub>: Intercept | -5.214 | 0.568 | -6.360 | -4.062 | 1.0009 | 6758.8 | 10022.2 | < 0.001 | *** |
|  | *β*<sub>0 50</sub>: Heat effect | 0.731 | 0.239 | 0.253 | 1.200 | 1.0000 | 11775.5 | 13060.2 | 0.004 | ** |
|  | *τ*<sub>*α* 50</sub>: RE SD Intercept | 1.453 | 0.498 | 0.762 | 2.684 | 1.0002 | 7700.6 | 10753.0 |  |  |
|  | *τ*<sub>*β* 50</sub>: RE SD Heat effect | 0.189 | 0.121 | 0.028 | 0.490 | 1.0000 | 12439.8 | 12656.5 |  |  |
|  | *ρ* (*τ*<sub>*α* 50</sub>, *τ*<sub>*β* 50</sub>): RE correlation | 0.028 | 0.553 | -0.931 | 0.939 | 1.0001 | 16632.8 | 13742.8 |  |  |
|  | *σ*<sub>50</sub>: Residual SD | 0.913 | 0.088 | 0.762 | 1.108 | 1.0003 | 7996.7 | 8026.7 |  |  |
| *P*<sub>88</sub> | *α*<sub>0 88</sub>: Intercept | -6.258 | 0.616 | -7.517 | -5.023 | 1.0003 | 7152.7 | 10214.5 | < 0.001 | *** |
|  | *β*<sub>0 88</sub>: Heat effect | 0.841 | 0.266 | 0.317 | 1.365 | 1.0000 | 12568.7 | 14116.0 | 0.003 | ** |
|  | *τ*<sub>*α* 88</sub>: RE SD Intercept | 1.603 | 0.532 | 0.863 | 2.919 | 1.0001 | 9182.8 | 10236.2 |  |  |
|  | *τ*<sub>*β* 88</sub>: RE SD Heat effect | 0.198 | 0.133 | 0.027 | 0.529 | 0.9999 | 11376.4 | 12809.2 |  |  |
|  | *ρ* (*τ*<sub>*α* 88</sub>, *τ*<sub>*β* 88</sub>): RE correlation | -0.174 | 0.565 | -0.972 | 0.908 | 1.0003 | 17688.6 | 14375.4 |  |  |
|  | *σ*<sub>88</sub>: Residual SD | 1.023 | 0.097 | 0.854 | 1.236 | 1.0003 | 10364.4 | 10659.3 |  |  |
| *P*<sub>ex</sub> | *α*<sub>0 *ex*</sub>: Intercept | -5.335 | 0.561 | -6.488 | -4.244 | 1.0004 | 5917.2 | 8491.5 | < 0.001 | *** |
|  | *β*<sub>0 *ex*</sub>: Heat effect | 0.746 | 0.208 | 0.331 | 1.152 | 1.0001 | 11070.1 | 12879.2 | < 0.001 | *** |
|  | *τ*<sub>*αex*</sub>: RE SD Intercept | 1.442 | 0.494 | 0.765 | 2.640 | 1.0004 | 7549.4 | 11051.5 |  |  |
|  | *τ*<sub>*βex*</sub>: RE SD Heat effect | 0.112 | 0.077 | 0.016 | 0.311 | 1.0001 | 10523.1 | 12607.7 |  |  |
|  | *ρ* (*τ*<sub>*αex*</sub>, *τ*<sub>*βex*</sub>): RE correlation | -0.194 | 0.607 | -0.984 | 0.940 | 1.0002 | 12785.6 | 13678.8 |  |  |
|  | *σ*<sub>*ex*</sub>: Residual SD | 0.828 | 0.079 | 0.693 | 1.001 | 1.0005 | 6868.1 | 7227.1 |  |  |
| Shared | *ρ* (*P*<sub>12</sub>, *P*<sub>50</sub>): resid. corr. | 0.771 | 0.056 | 0.647 | 0.863 | 1.0004 | 8718.0 | 10861.6 |  |  |
|  | *ρ* (*P*<sub>12</sub>, *P*<sub>88</sub>): resid. corr. | 0.537 | 0.095 | 0.333 | 0.704 | 1.0004 | 9190.1 | 10633.2 |  |  |
|  | *ρ* (*P*<sub>50</sub>, *P*<sub>88</sub>): resid. corr. | 0.754 | 0.059 | 0.621 | 0.853 | 1.0002 | 10452.5 | 12086.0 |  |  |
|  | *ρ* (*P*<sub>12</sub>, *P*<sub>*ex*</sub>): resid. corr. | 0.827 | 0.043 | 0.731 | 0.897 | 1.0006 | 8537.0 | 9934.6 |  |  |
|  | *ρ* (*P*<sub>50</sub>, *P*<sub>*ex*</sub>): resid. corr. | 0.938 | 0.017 | 0.899 | 0.965 | 1.0004 | 9762.9 | 10839.2 |  |  |
|  | *ρ* (*P*<sub>88</sub>, *P*<sub>*ex*</sub>): resid. corr. | 0.882 | 0.030 | 0.813 | 0.931 | 1.0001 | 12589.7 | 12998.0 |  |  |

**Table S2.**
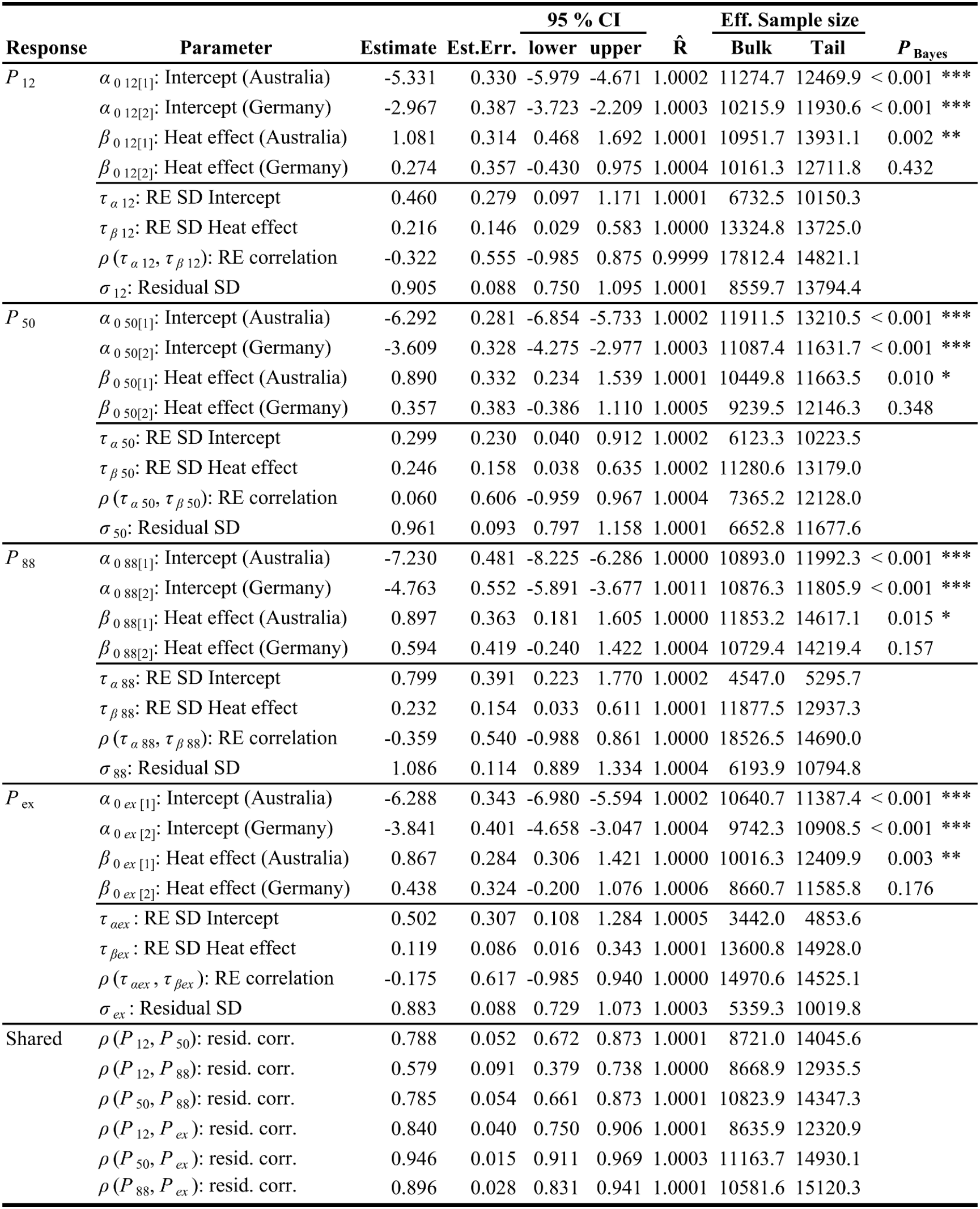
Results of the multi-response mixed model of *P*_12_, *P*_50_, *P*_88_, and *P*_ex_ with separate parameters for Australia and Germany. See Notes S1.2 for model structure. Shown are the posterior means of the parameter estimates with their MCMC standard error, 95 % credible interval, potential scale reduction factor 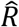, bulk and tail effective sample sizes and a Bayesian pseudo-p-value (only shown for fixed effects, see Notes S1.4). Symbols indicate parameters credibly different from zero on the 0.1 (.), 0.05 (*), 0.01 (**) and 0.001 level (***).

|  |  |  |  | 95 % CI |  |  | Eff. Sample size |  |  |  |
| Response | Parameter | Estimate | Est.Err. | lower | upper | R̂ | Bulk | Tail | *P*<sub>Bayes</sub> |  |
|---|---|---|---|---|---|---|---|---|---|---|
| *P*<sub>12</sub> | *α*<sub>0 12[1]</sub>: Intercept (Australia) | -5.331 | 0.330 | -5.979 | -4.671 | 1.0002 | 11274.7 | 12469.9 | < 0.001 | *** |
|  | *α*<sub>0 12[2]</sub>: Intercept (Germany) | -2.967 | 0.387 | -3.723 | -2.209 | 1.0003 | 10215.9 | 11930.6 | < 0.001 | *** |
|  | *β*<sub>0 12[1]</sub>: Heat effect (Australia) | 1.081 | 0.314 | 0.468 | 1.692 | 1.0001 | 10951.7 | 13931.1 | 0.002 | ** |
|  | *β*<sub>0 12[2]</sub>: Heat effect (Germany) | 0.274 | 0.357 | -0.430 | 0.975 | 1.0004 | 10161.3 | 12711.8 | 0.432 |  |
|  | *τ*<sub>*α* 12</sub>: RE SD Intercept | 0.460 | 0.279 | 0.097 | 1.171 | 1.0001 | 6732.5 | 10150.3 |  |  |
|  | *τ*<sub>*β* 12</sub>: RE SD Heat effect | 0.216 | 0.146 | 0.029 | 0.583 | 1.0000 | 13324.8 | 13725.0 |  |  |
|  | *ρ* (*τ*<sub>*α* 12</sub>, *τ*<sub>*β* 12</sub>): RE correlation | -0.322 | 0.555 | -0.985 | 0.875 | 0.9999 | 17812.4 | 14821.1 |  |  |
|  | *σ*<sub>12</sub>: Residual SD | 0.905 | 0.088 | 0.750 | 1.095 | 1.0001 | 8559.7 | 13794.4 |  |  |
| *P*<sub>50</sub> | *α*<sub>0 50[1]</sub>: Intercept (Australia) | -6.292 | 0.281 | -6.854 | -5.733 | 1.0002 | 11911.5 | 13210.5 | < 0.001 | *** |
|  | *α*<sub>0 50[2]</sub>: Intercept (Germany) | -3.609 | 0.328 | -4.275 | -2.977 | 1.0003 | 11087.4 | 11631.7 | < 0.001 | *** |
|  | *β*<sub>0 50[1]</sub>: Heat effect (Australia) | 0.890 | 0.332 | 0.234 | 1.539 | 1.0001 | 10449.8 | 11663.5 | 0.010 | * |
|  | *β*<sub>0 50[2]</sub>: Heat effect (Germany) | 0.357 | 0.383 | -0.386 | 1.110 | 1.0005 | 9239.5 | 12146.3 | 0.348 |  |
|  | *τ*<sub>*α* 50</sub>: RE SD Intercept | 0.299 | 0.230 | 0.040 | 0.912 | 1.0002 | 6123.3 | 10223.5 |  |  |
|  | *τ*<sub>*β* 50</sub>: RE SD Heat effect | 0.246 | 0.158 | 0.038 | 0.635 | 1.0002 | 11280.6 | 13179.0 |  |  |
|  | *ρ* (*τ*<sub>*α* 50</sub>, *τ*<sub>*β* 50</sub>): RE correlation | 0.060 | 0.606 | -0.959 | 0.967 | 1.0004 | 7365.2 | 12128.0 |  |  |
|  | *σ*<sub>50</sub>: Residual SD | 0.961 | 0.093 | 0.797 | 1.158 | 1.0001 | 6652.8 | 11677.6 |  |  |
| *P*<sub>88</sub> | *α*<sub>0 88[1]</sub>: Intercept (Australia) | -7.230 | 0.481 | -8.225 | -6.286 | 1.0000 | 10893.0 | 11992.3 | < 0.001 | *** |
|  | *α*<sub>0 88[2]</sub>: Intercept (Germany) | -4.763 | 0.552 | -5.891 | -3.677 | 1.0011 | 10876.3 | 11805.9 | < 0.001 | *** |
|  | *β*<sub>0 88[1]</sub>: Heat effect (Australia) | 0.897 | 0.363 | 0.181 | 1.605 | 1.0000 | 11853.2 | 14617.1 | 0.015 | * |
|  | *β*<sub>0 88[2]</sub>: Heat effect (Germany) | 0.594 | 0.419 | -0.240 | 1.422 | 1.0004 | 10729.4 | 14219.4 | 0.157 |  |
|  | *τ*<sub>*α* 88</sub>: RE SD Intercept | 0.799 | 0.391 | 0.223 | 1.770 | 1.0002 | 4547.0 | 5295.7 |  |  |
|  | *τ*<sub>*β* 88</sub>: RE SD Heat effect | 0.232 | 0.154 | 0.033 | 0.611 | 1.0001 | 11877.5 | 12937.3 |  |  |
|  | *ρ* (*τ*<sub>*α* 88</sub>, *τ*<sub>*β* 88</sub>): RE correlation | -0.359 | 0.540 | -0.988 | 0.861 | 1.0000 | 18526.5 | 14690.0 |  |  |
|  | *σ*<sub>88</sub>: Residual SD | 1.086 | 0.114 | 0.889 | 1.334 | 1.0004 | 6193.9 | 10794.8 |  |  |
| *P*<sub>ex</sub> | *α*<sub>0 *ex* [1]</sub>: Intercept (Australia) | -6.288 | 0.343 | -6.980 | -5.594 | 1.0002 | 10640.7 | 11387.4 | < 0.001 | *** |
|  | *α*<sub>0 *ex* [2]</sub>: Intercept (Germany) | -3.841 | 0.401 | -4.658 | -3.047 | 1.0004 | 9742.3 | 10908.5 | < 0.001 | *** |
|  | *β*<sub>0 *ex* [1]</sub>: Heat effect (Australia) | 0.867 | 0.284 | 0.306 | 1.421 | 1.0000 | 10016.3 | 12409.9 | 0.003 | ** |
|  | *β*<sub>0 *ex* [2]</sub>: Heat effect (Germany) | 0.438 | 0.324 | -0.200 | 1.076 | 1.0006 | 8660.7 | 11585.8 | 0.176 |  |
|  | *τ*<sub>*αex*</sub>: RE SD Intercept | 0.502 | 0.307 | 0.108 | 1.284 | 1.0005 | 3442.0 | 4853.6 |  |  |
|  | *τ*<sub>*βex*</sub>: RE SD Heat effect | 0.119 | 0.086 | 0.016 | 0.343 | 1.0001 | 13600.8 | 14928.0 |  |  |
|  | *ρ* (*τ*<sub>*αex*</sub>, *τ*<sub>*βex*</sub>): RE correlation | -0.175 | 0.617 | -0.985 | 0.940 | 1.0000 | 14970.6 | 14525.1 |  |  |
|  | *σ*<sub>*ex*</sub>: Residual SD | 0.883 | 0.088 | 0.729 | 1.073 | 1.0003 | 5359.3 | 10019.8 |  |  |
| Shared | *ρ* (*P*<sub>12</sub>, *P*<sub>50</sub>): resid. corr. | 0.788 | 0.052 | 0.672 | 0.873 | 1.0001 | 8721.0 | 14045.6 |  |  |
|  | *ρ* (*P*<sub>12</sub>, *P*<sub>88</sub>): resid. corr. | 0.579 | 0.091 | 0.379 | 0.738 | 1.0000 | 8668.9 | 12935.5 |  |  |
|  | *ρ* (*P*<sub>50</sub>, *P*<sub>88</sub>): resid. corr. | 0.785 | 0.054 | 0.661 | 0.873 | 1.0001 | 10823.9 | 14347.3 |  |  |
|  | *ρ* (*P*<sub>12</sub>, *P*<sub>*ex*</sub>): resid. corr. | 0.840 | 0.040 | 0.750 | 0.906 | 1.0001 | 8635.9 | 12320.9 |  |  |
|  | *ρ* (*P*<sub>50</sub>, *P*<sub>*ex*</sub>): resid. corr. | 0.946 | 0.015 | 0.911 | 0.969 | 1.0003 | 11163.7 | 14930.1 |  |  |
|  | *ρ* (*P*<sub>88</sub>, *P*<sub>*ex*</sub>): resid. corr. | 0.896 | 0.028 | 0.831 | 0.941 | 1.0001 | 10581.6 | 15120.3 |  |  |

**Table S3.**
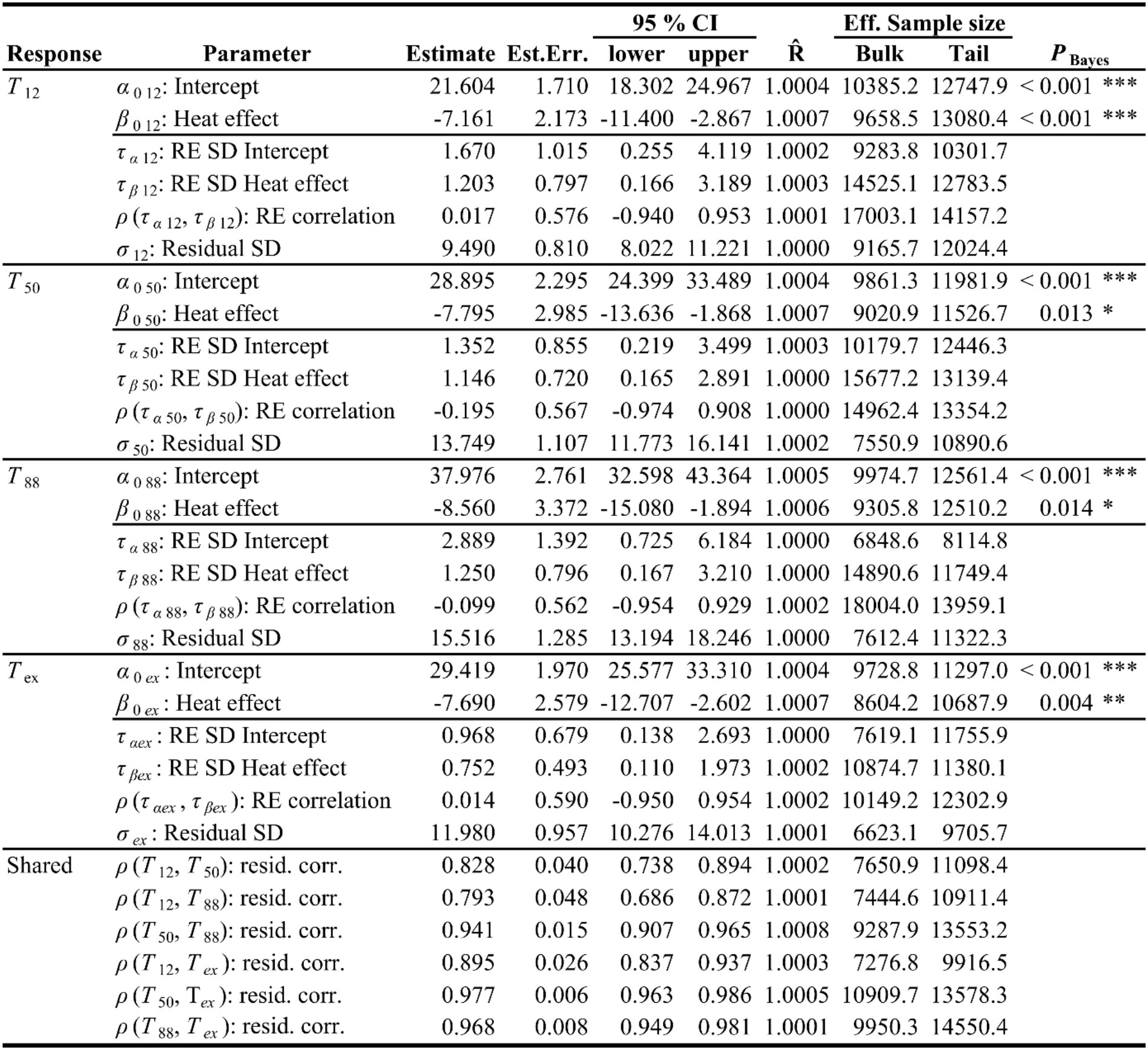
Results of the multi-response mixed model of *T*_12_, *T*_50_, *T*_88_, and *T*_ex_ pooled over both locations. See Notes S1.2 for model structure. Shown are the posterior means of the parameter estimates with their MCMC standard error, 95 % credible interval, potential scale reduction factor 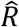, bulk and tail effective sample sizes and a Bayesian pseudo-p-value (only shown for fixed effects, see Notes S1.4). Symbols indicate parameters credibly different from zero on the 0.1 (.), 0.05 (*), 0.01 (**) and 0.001 level (***).

|  |  |  |  | 95 % CI |  |  | Eff. Sample size |  |  |
|---|---|---|---|---|---|---|---|---|---|
| Response | Parameter | Estimate | Est.Err. | lower | upper | R̂ | Bulk | Tail | ***P***<sub>**Bayes**</sub> |
| *T*<sub>12</sub> | *α*<sub>0 12</sub>: Intercept | 21.604 | 1.710 | 18.302 | 24.967 | 1.0004 | 10385.2 | 12747.9 | < 0.001 *** |
|  | *β*<sub>0 12</sub>: Heat effect | -7.161 | 2.173 | -11.400 | -2.867 | 1.0007 | 9658.5 | 13080.4 | < 0.001 *** |
|  | *τ*<sub>*α* 12</sub>: RE SD Intercept | 1.670 | 1.015 | 0.255 | 4.119 | 1.0002 | 9283.8 | 10301.7 |  |
|  | *τ*<sub>*β* 12</sub>: RE SD Heat effect | 1.203 | 0.797 | 0.166 | 3.189 | 1.0003 | 14525.1 | 12783.5 |  |
|  | *ρ* (*τ*<sub>*α* 12</sub>, *τ*<sub>*β* 12</sub>): RE correlation | 0.017 | 0.576 | -0.940 | 0.953 | 1.0001 | 17003.1 | 14157.2 |  |
|  | *σ*<sub>12</sub>: Residual SD | 9.490 | 0.810 | 8.022 | 11.221 | 1.0000 | 9165.7 | 12024.4 |  |
| *T*<sub>50</sub> | *α*<sub>0 50</sub>: Intercept | 28.895 | 2.295 | 24.399 | 33.489 | 1.0004 | 9861.3 | 11981.9 | < 0.001 *** |
|  | *β*<sub>0 50</sub>: Heat effect | -7.795 | 2.985 | -13.636 | -1.868 | 1.0007 | 9020.9 | 11526.7 | 0.013 * |
|  | *τ*<sub>*α* 50</sub>: RE SD Intercept | 1.352 | 0.855 | 0.219 | 3.499 | 1.0003 | 10179.7 | 12446.3 |  |
|  | *τ*<sub>*β* 50</sub>: RE SD Heat effect | 1.146 | 0.720 | 0.165 | 2.891 | 1.0000 | 15677.2 | 13139.4 |  |
|  | *ρ* (*τ*<sub>*α* 50</sub>, *τ*<sub>*β* 50</sub>): RE correlation | -0.195 | 0.567 | -0.974 | 0.908 | 1.0000 | 14962.4 | 13354.2 |  |
|  | *σ*<sub>50</sub>: Residual SD | 13.749 | 1.107 | 11.773 | 16.141 | 1.0002 | 7550.9 | 10890.6 |  |
| *T*<sub>88</sub> | *α*<sub>0 88</sub>: Intercept | 37.976 | 2.761 | 32.598 | 43.364 | 1.0005 | 9974.7 | 12561.4 | < 0.001 *** |
|  | *β*<sub>0 88</sub>: Heat effect | -8.560 | 3.372 | -15.080 | -1.894 | 1.0006 | 9305.8 | 12510.2 | 0.014 * |
|  | *τ*<sub>*α* 88</sub>: RE SD Intercept | 2.889 | 1.392 | 0.725 | 6.184 | 1.0000 | 6848.6 | 8114.8 |  |
|  | *τ*<sub>*β* 88</sub>: RE SD Heat effect | 1.250 | 0.796 | 0.167 | 3.210 | 1.0000 | 14890.6 | 11749.4 |  |
|  | *ρ* (*τ*<sub>*α* 88</sub>, *τ*<sub>*β* 88</sub>): RE correlation | -0.099 | 0.562 | -0.954 | 0.929 | 1.0002 | 18004.0 | 13959.1 |  |
|  | *σ*<sub>88</sub>: Residual SD | 15.516 | 1.285 | 13.194 | 18.246 | 1.0000 | 7612.4 | 11322.3 |  |
| *T*<sub>ex</sub> | *α*<sub>0 *ex*</sub>: Intercept | 29.419 | 1.970 | 25.577 | 33.310 | 1.0004 | 9728.8 | 11297.0 | < 0.001 *** |
|  | *β*<sub>0 *ex*</sub>: Heat effect | -7.690 | 2.579 | -12.707 | -2.602 | 1.0007 | 8604.2 | 10687.9 | 0.004 ** |
|  | *τ*<sub>*αex*</sub>: RE SD Intercept | 0.968 | 0.679 | 0.138 | 2.693 | 1.0000 | 7619.1 | 11755.9 |  |
|  | *τ*<sub>*βex*</sub>: RE SD Heat effect | 0.752 | 0.493 | 0.110 | 1.973 | 1.0002 | 10874.7 | 11380.1 |  |
|  | *ρ* (*τ*<sub>*αex*</sub>, *τ*<sub>*βex*</sub>): RE correlation | 0.014 | 0.590 | -0.950 | 0.954 | 1.0002 | 10149.2 | 12302.9 |  |
|  | *σ*<sub>*ex*</sub>: Residual SD | 11.980 | 0.957 | 10.276 | 14.013 | 1.0001 | 6623.1 | 9705.7 |  |
| Shared | *ρ* (*T*<sub>12</sub>, *T*<sub>50</sub>): resid. corr. | 0.828 | 0.040 | 0.738 | 0.894 | 1.0002 | 7650.9 | 11098.4 |  |
|  | *ρ* (*T*<sub>12</sub>, *T*<sub>88</sub>): resid. corr. | 0.793 | 0.048 | 0.686 | 0.872 | 1.0001 | 7444.6 | 10911.4 |  |
|  | *ρ* (*T*<sub>50</sub>, *T*<sub>88</sub>): resid. corr. | 0.941 | 0.015 | 0.907 | 0.965 | 1.0008 | 9287.9 | 13553.2 |  |
|  | *ρ* (*T*<sub>12</sub>, *T*<sub>*ex*</sub>): resid. corr. | 0.895 | 0.026 | 0.837 | 0.937 | 1.0003 | 7276.8 | 9916.5 |  |
|  | *ρ* (*T*<sub>50</sub>, T<sub>*ex*</sub>): resid. corr. | 0.977 | 0.006 | 0.963 | 0.986 | 1.0005 | 10909.7 | 13578.3 |  |
|  | *ρ* (*T*<sub>88</sub>, *T*<sub>*ex*</sub>): resid. corr. | 0.968 | 0.008 | 0.949 | 0.981 | 1.0001 | 9950.3 | 14550.4 |  |

**Table S4.**
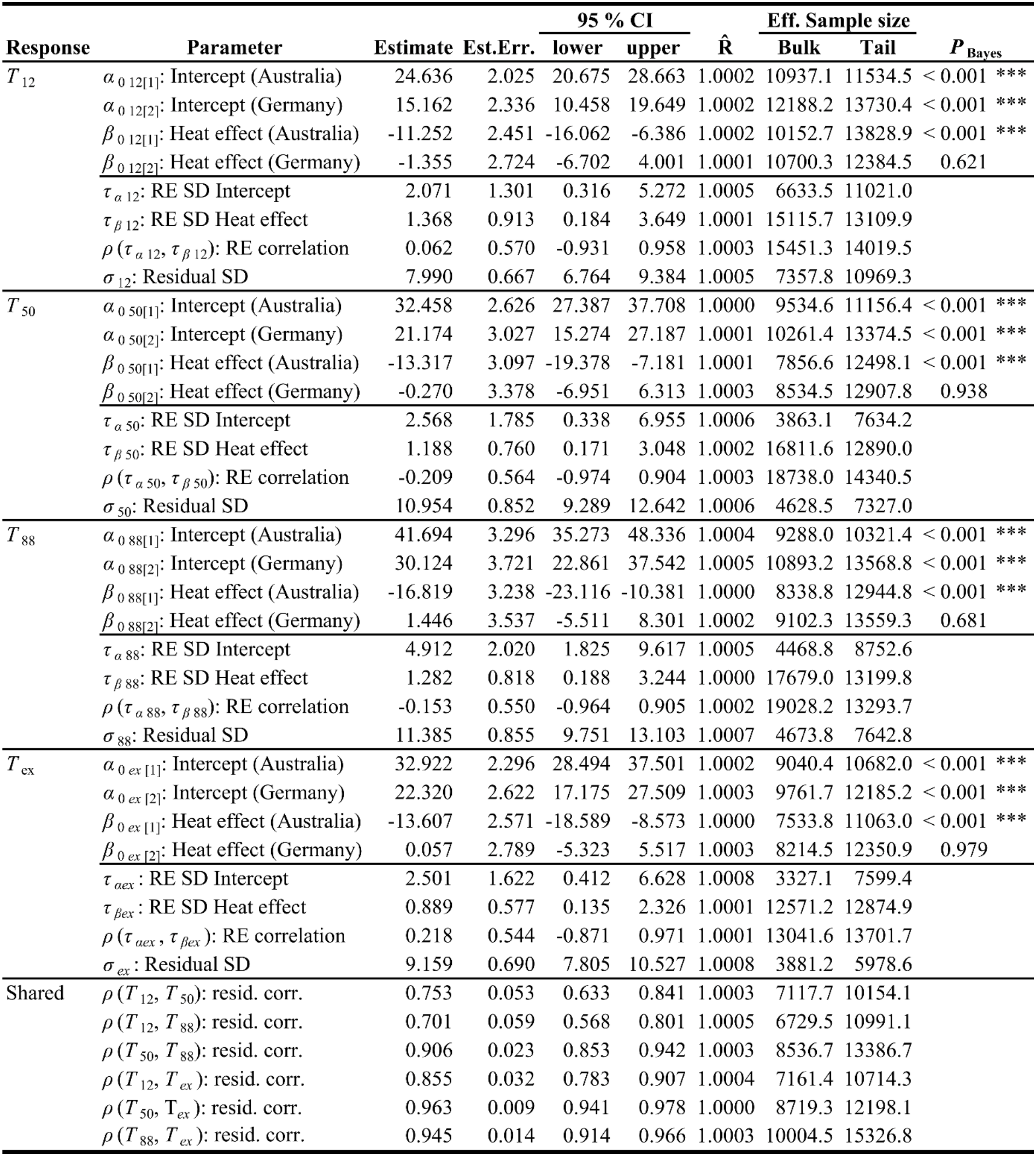
Results of the multi-response mixed model of *T*_12_, *T*_50_, *T*_88_, and *T*_ex_ with separate parameters for Australia and Germany. See Notes S1.2 for model structure. Shown are the posterior means of the parameter estimates with their MCMC standard error, 95 % credible interval, potential scale reduction factor 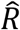, bulk and tail effective sample sizes and a Bayesian pseudo-p-value (only shown for fixed effects, see Notes S1.4). Symbols indicate parameters credibly different from zero on the 0.1 (.), 0.05 (*), 0.01 (**) and 0.001 level (***).

|  |  |  |  | 95 % CI |  |  | Eff. Sample size |  |  |  |
| Response | Parameter | Estimate | Est.Err. | lower | upper | R̂ | Bulk | Tail | *P*<sub>Bayes</sub> |  |
|---|---|---|---|---|---|---|---|---|---|---|
| *T*<sub>12</sub> | α<sub>0 12[1]</sub>: Intercept (Australia) | 24.636 | 2.025 | 20.675 | 28.663 | 1.0002 | 10937.1 | 11534.5 | < 0.001 | *** |
|  | α<sub>0 12[2]</sub>: Intercept (Germany) | 15.162 | 2.336 | 10.458 | 19.649 | 1.0002 | 12188.2 | 13730.4 | < 0.001 | *** |
|  | β<sub>0 12[1]</sub>: Heat effect (Australia) | -11.252 | 2.451 | -16.062 | -6.386 | 1.0002 | 10152.7 | 13828.9 | < 0.001 | *** |
|  | β<sub>0 12[2]</sub>: Heat effect (Germany) | -1.355 | 2.724 | -6.702 | 4.001 | 1.0001 | 10700.3 | 12384.5 | 0.621 |  |
|  | τ<sub>α 12</sub>: RE SD Intercept | 2.071 | 1.301 | 0.316 | 5.272 | 1.0005 | 6633.5 | 11021.0 |  |  |
|  | τ<sub>β 12</sub>: RE SD Heat effect | 1.368 | 0.913 | 0.184 | 3.649 | 1.0001 | 15115.7 | 13109.9 |  |  |
|  | ρ (τ<sub>α 12</sub>, τ<sub>β 12</sub>): RE correlation | 0.062 | 0.570 | -0.931 | 0.958 | 1.0003 | 15451.3 | 14019.5 |  |  |
|  | σ<sub>12</sub>: Residual SD | 7.990 | 0.667 | 6.764 | 9.384 | 1.0005 | 7357.8 | 10969.3 |  |  |
| *T*<sub>50</sub> | α<sub>0 50[1]</sub>: Intercept (Australia) | 32.458 | 2.626 | 27.387 | 37.708 | 1.0000 | 9534.6 | 11156.4 | < 0.001 | *** |
|  | α<sub>0 50[2]</sub>: Intercept (Germany) | 21.174 | 3.027 | 15.274 | 27.187 | 1.0001 | 10261.4 | 13374.5 | < 0.001 | *** |
|  | β<sub>0 50[1]</sub>: Heat effect (Australia) | -13.317 | 3.097 | -19.378 | -7.181 | 1.0001 | 7856.6 | 12498.1 | < 0.001 | *** |
|  | β<sub>0 50[2]</sub>: Heat effect (Germany) | -0.270 | 3.378 | -6.951 | 6.313 | 1.0003 | 8534.5 | 12907.8 | 0.938 |  |
|  | τ<sub>α 50</sub>: RE SD Intercept | 2.568 | 1.785 | 0.338 | 6.955 | 1.0006 | 3863.1 | 7634.2 |  |  |
|  | τ<sub>β 50</sub>: RE SD Heat effect | 1.188 | 0.760 | 0.171 | 3.048 | 1.0002 | 16811.6 | 12890.0 |  |  |
|  | ρ (τ<sub>α 50</sub>, τ<sub>β 50</sub>): RE correlation | -0.209 | 0.564 | -0.974 | 0.904 | 1.0003 | 18738.0 | 14340.5 |  |  |
|  | σ<sub>50</sub>: Residual SD | 10.954 | 0.852 | 9.289 | 12.642 | 1.0006 | 4628.5 | 7327.0 |  |  |
| *T*<sub>88</sub> | α<sub>0 88[1]</sub>: Intercept (Australia) | 41.694 | 3.296 | 35.273 | 48.336 | 1.0004 | 9288.0 | 10321.4 | < 0.001 | *** |
|  | α<sub>0 88[2]</sub>: Intercept (Germany) | 30.124 | 3.721 | 22.861 | 37.542 | 1.0005 | 10893.2 | 13568.8 | < 0.001 | *** |
|  | β<sub>0 88[1]</sub>: Heat effect (Australia) | -16.819 | 3.238 | -23.116 | -10.381 | 1.0000 | 8338.8 | 12944.8 | < 0.001 | *** |
|  | β<sub>0 88[2]</sub>: Heat effect (Germany) | 1.446 | 3.537 | -5.511 | 8.301 | 1.0002 | 9102.3 | 13559.3 | 0.681 |  |
|  | τ<sub>α 88</sub>: RE SD Intercept | 4.912 | 2.020 | 1.825 | 9.617 | 1.0005 | 4468.8 | 8752.6 |  |  |
|  | τ<sub>β 88</sub>: RE SD Heat effect | 1.282 | 0.818 | 0.188 | 3.244 | 1.0000 | 17679.0 | 13199.8 |  |  |
|  | ρ (τ<sub>α 88</sub>, τ<sub>β 88</sub>): RE correlation | -0.153 | 0.550 | -0.964 | 0.905 | 1.0002 | 19028.2 | 13293.7 |  |  |
|  | σ<sub>88</sub>: Residual SD | 11.385 | 0.855 | 9.751 | 13.103 | 1.0007 | 4673.8 | 7642.8 |  |  |
| *T*<sub>ex</sub> | α<sub>0 *ex* [1]</sub>: Intercept (Australia) | 32.922 | 2.296 | 28.494 | 37.501 | 1.0002 | 9040.4 | 10682.0 | < 0.001 | *** |
|  | α<sub>0 *ex* [2]</sub>: Intercept (Germany) | 22.320 | 2.622 | 17.175 | 27.509 | 1.0003 | 9761.7 | 12185.2 | < 0.001 | *** |
|  | β<sub>0 *ex* [1]</sub>: Heat effect (Australia) | -13.607 | 2.571 | -18.589 | -8.573 | 1.0000 | 7533.8 | 11063.0 | < 0.001 | *** |
|  | β<sub>0 *ex* [2]</sub>: Heat effect (Germany) | 0.057 | 2.789 | -5.323 | 5.517 | 1.0003 | 8214.5 | 12350.9 | 0.979 |  |
|  | τ<sub>α*ex*</sub>: RE SD Intercept | 2.501 | 1.622 | 0.412 | 6.628 | 1.0008 | 3327.1 | 7599.4 |  |  |
|  | τ<sub>β*ex*</sub>: RE SD Heat effect | 0.889 | 0.577 | 0.135 | 2.326 | 1.0001 | 12571.2 | 12874.9 |  |  |
|  | ρ (τ<sub>α*ex*</sub>, τ<sub>β*ex*</sub>): RE correlation | 0.218 | 0.544 | -0.871 | 0.971 | 1.0001 | 13041.6 | 13701.7 |  |  |
|  | σ<sub>*ex*</sub>: Residual SD | 9.159 | 0.690 | 7.805 | 10.527 | 1.0008 | 3881.2 | 5978.6 |  |  |
| Shared | ρ (*T*<sub>12</sub>, *T*<sub>50</sub>): resid. corr. | 0.753 | 0.053 | 0.633 | 0.841 | 1.0003 | 7117.7 | 10154.1 |  |  |
|  | ρ (*T*<sub>12</sub>, *T*<sub>88</sub>): resid. corr. | 0.701 | 0.059 | 0.568 | 0.801 | 1.0005 | 6729.5 | 10991.1 |  |  |
|  | ρ (*T*<sub>50</sub>, *T*<sub>88</sub>): resid. corr. | 0.906 | 0.023 | 0.853 | 0.942 | 1.0003 | 8536.7 | 13386.7 |  |  |
|  | ρ (*T*<sub>12</sub>, *T*<sub>*ex*</sub>): resid. corr. | 0.855 | 0.032 | 0.783 | 0.907 | 1.0004 | 7161.4 | 10714.3 |  |  |
|  | ρ (*T*<sub>50</sub>, T<sub>*ex*</sub>): resid. corr. | 0.963 | 0.009 | 0.941 | 0.978 | 1.0000 | 8719.3 | 12198.1 |  |  |
|  | ρ (*T*<sub>88</sub>, *T*<sub>*ex*</sub>): resid. corr. | 0.945 | 0.014 | 0.914 | 0.966 | 1.0003 | 10004.5 | 15326.8 |  |  |

